# Individualized Gabor Flicker for Well-Tolerated Neural Entrainment During Working Memory

**DOI:** 10.64898/2026.10.01.755870

**Authors:** Clément Blanc, Frédéric Dehais, Sébastien Scannella

**Affiliations:** Fédération ENAC ISAE-SUPAERO ONERA, Université de Toulouse, 10 Avenue Marc Pélegrin, Toulouse

**Keywords:** Neuromodulation, Steady-state visual evoked potentials (SSVEPs), Visual entrainment, Working memory, Gabor flicker

## Abstract

Rhythmic visual stimulation offers a non-invasive means of modulating oscillatory brain activity, but conventional flicker may induce considerable visual discomfort. We developed an individualized Gabor flicker stimulation designed to balance neural entrainment and tolerability. In Experiment 1, thirty-four participants passively viewed Plain and Gabor flickers presented at their individual alpha frequency (IAF), IAF *−*2 Hz, or IAF +2 Hz, and at threshold-adjusted or full intensity. Steady-state visual evoked potential signal-to-noise ratio (SNR) was highest at IAF and increased with intensity. Plain and Gabor flickers produced comparable overall SNR, but Gabor flicker was rated as substantially more comfortable and less mentally fatiguing, with stimulation at IAF providing the most favorable balance between neural entrainment and tolerability. In Experiment 2, the optimized stimulation parameters were applied during a retro-cue working memory task using individualized theta and alpha frequency flickers. Stimulation increased SNR and inter-trial phase coherence relative to a non rhythmic Natural condition. No behavioral benefit was observed in the full sample. However, an exploratory subgroup analysis indicated higher accuracy with theta stimulation of cued representations and alpha stimulation of uncued representations than in the Natural condition among participants who showed limited baseline retro-cue benefits. These findings show that individualized Gabor flicker can provide robust and well-tolerated neural entrainment during passive viewing and working memory retention, although its behavioral effects may vary across individuals according to their cognitive performance.

## 1 Introduction

Non-invasive rhythmic stimulation provides a means of probing and potentially modulating oscillatory mechanisms involved in cognition. Neural oscillations contribute to the temporal coordination of attention, memory, and cognitive control (Buzśaki & Draguhn, 2004; Fries, 2015), and frequency-specific stimulation has therefore been used to investigate their causal role in behavior (De Graaf et al., 2013; Riddle et al., 2020). Among available approaches, rhythmic visual stimulation is particularly attractive because it is non-contact, requires minimal specialized equipment, and reliably induces frequency-specific neural responses in the form of steady-state visual evoked potentials (SSVEPs) (Norcia et al., 2015; Vialatte et al., 2010). Importantly, visual rhythmic stimulation can also influence cognition: alpha-frequency stimulation has been shown to modulate selective attention (Gulbinaite et al., 2017), whereas theta-frequency stimulation has been associated with changes in working-memory performance (Pileckyte & Soto-Faraco, 2024).

The efficacy of rhythmic stimulation may depend critically on the relationship between the stimulation frequency and the intrinsic oscillatory dynamics of the individual. Endogenous oscillatory peaks vary across participants, particularly within the alpha range, such that a fixed stimulation frequency may closely match the dominant rhythm of one participant while being substantially detuned in another. Consistent with this view, Gulbinaite and colleagues showed that the behavioral effects of 10-Hz visual stimulation depended on its proximity to the participant’s individual alpha frequency (IAF) (Gulbinaite et al., 2017). Neural responses to rhythmic visual stimulation also exhibit resonance-like properties, with stronger responses when external stimulation approaches endogenous oscillatory frequencies (Herrmann, 2001; Notbohm & Herrmann, 2016). These findings support the use of individualized rather than fixed-frequency stimulation protocols.

A major limitation of visual neuromodulation, however, is the perceptual burden associated with conventional stimuli. Salient luminance modulation, uniform flashes, and high-contrast patterns can efficiently drive neural responses but are also prone to produce visual discomfort, eye strain, distraction, and mental fatigue during prolonged exposure (Fisher et al., 2005). This is particularly problematic in cognitive paradigms, where stimulation must often be sustained and repeatedly delivered over extended experimental sessions. Visual discomfort may consequently reduce task engagement and confound the behavioral effects attributed to stimulation.

To address this limitation, we recently developed spatially textured visual stimulation designed to preserve frequency-specific neural responses while reducing the perceptual salience of conventional flicker. Previous work using Gabor-and Ricker-like textures showed substantially improved subjective comfort while maintaining robust visually evoked responses and supporting alpha-band neuromodulation (Blanc et al., 2026; Dehais et al., 2024). Such stimuli may therefore provide a better compromise between neural efficacy and tolerability, making them particularly suitable for sustained cognitive neuromodulation.

Two questions nevertheless remain to be addressed. First, it is unclear whether the improved tolerability of textured stimulation can be combined with individualized frequency targeting without compromising neural responsiveness. Stimulus structure, intensity, and the match between stimulation frequency and endogenous rhythms may all influence the resulting neural response. We therefore first tested the general principle that stimulation delivered near an individual’s endogenous oscillatory frequency may enhance neural responsiveness. We focused on the alpha band because the occipital alpha peak can be robustly estimated from resting-state EEG. This optimization stage then provided the basis for extending individualized frequency stimulation to both alpha and theta bands in a working-memory experiment.

Second, once these parameters have been optimized, it remains to be determined whether individualized textured stimulation can be used to target oscillatory processes involved in working memory. It has been shown that working-memory prioritization partly relies on dissociable theta- and alpha-band mechanisms. While theta activity has been associated with active maintenance, executive coordination, and frontoparietal communication (Cavanagh & Frank, 2014; Sauseng et al., 2005; Wolinski et al., 2018), posterior alpha activity has been linked to attentional inhibition and the down-weighting of irrelevant representations (Jensen & Mazaheri, 2010; Sauseng et al., 2009). In a retro-cue paradigm, Riddle and colleagues demonstrated frequency-specific contributions of these rhythms to prioritized and non-prioritized memory representations (Riddle et al., 2020). This paradigm therefore provides a suitable framework for evaluating whether individualized rhythmic visual stimulation can engage comparable mechanisms using a more tolerable and readily deployable stimulation method.

The present study addressed these questions in two successive experiments. Experiment 1 served as an optimization stage and tested whether visually comfortable Gabor stimulation could be combined with individualized frequency targeting while maintaining robust neural responses. During passive viewing, we manipulated stimulation frequency relative to each participant’s IAF (IAF *vs.* IAF *−*2 Hz *vs.* IAF +2 Hz), stimulus format (Plain *vs.* Gabor flicker), and intensity (Full *vs.* Threshold). SSVEP responses, visual comfort, and mental fatigue were assessed to identify the stimulation configuration providing the most favorable balance between neural efficacy and tolerability.

Experiment 2 was then developed by applying the optimized stimulation parameters during a retro-cue working-memory task. Individualized theta- and alpha-frequency Gabor flickers were presented during memory retention to test whether frequency-specific stimulation of prioritized and non-prioritized representations could modulate neural activity and behavioral performance.

## 2 EXPERIMENT 1

### 2.1 Materials and Methods

#### 2.1.1 Participants

Thirty-four volunteers participated in the study (mean age = 24.5 years, SD = 3.74; 21 males). All participants had normal or corrected-to-normal vision. None reported neurological, psychological, or psychiatric disorders. All provided their written informed consent to participate in this study that was approved by the local Research Ethics Committee of the Federal University of Toulouse (CER No. 2024-857), and conformed to the principles expressed in the Declaration of Helsinki. Participants were compensated €10 for participating in the study.

#### 2.1.2 Stimuli and procedure

The experiment followed a fully within-subject 3 *×* 2 *×* 2 design manipulating stimulation frequency (IAF *vs.* IAF *−*2 Hz *vs.* IAF +2 Hz), flicker type (Plain *vs.* Gabor; Figure 1), and intensity (Full *vs.* Threshold).

**Figure 1:**
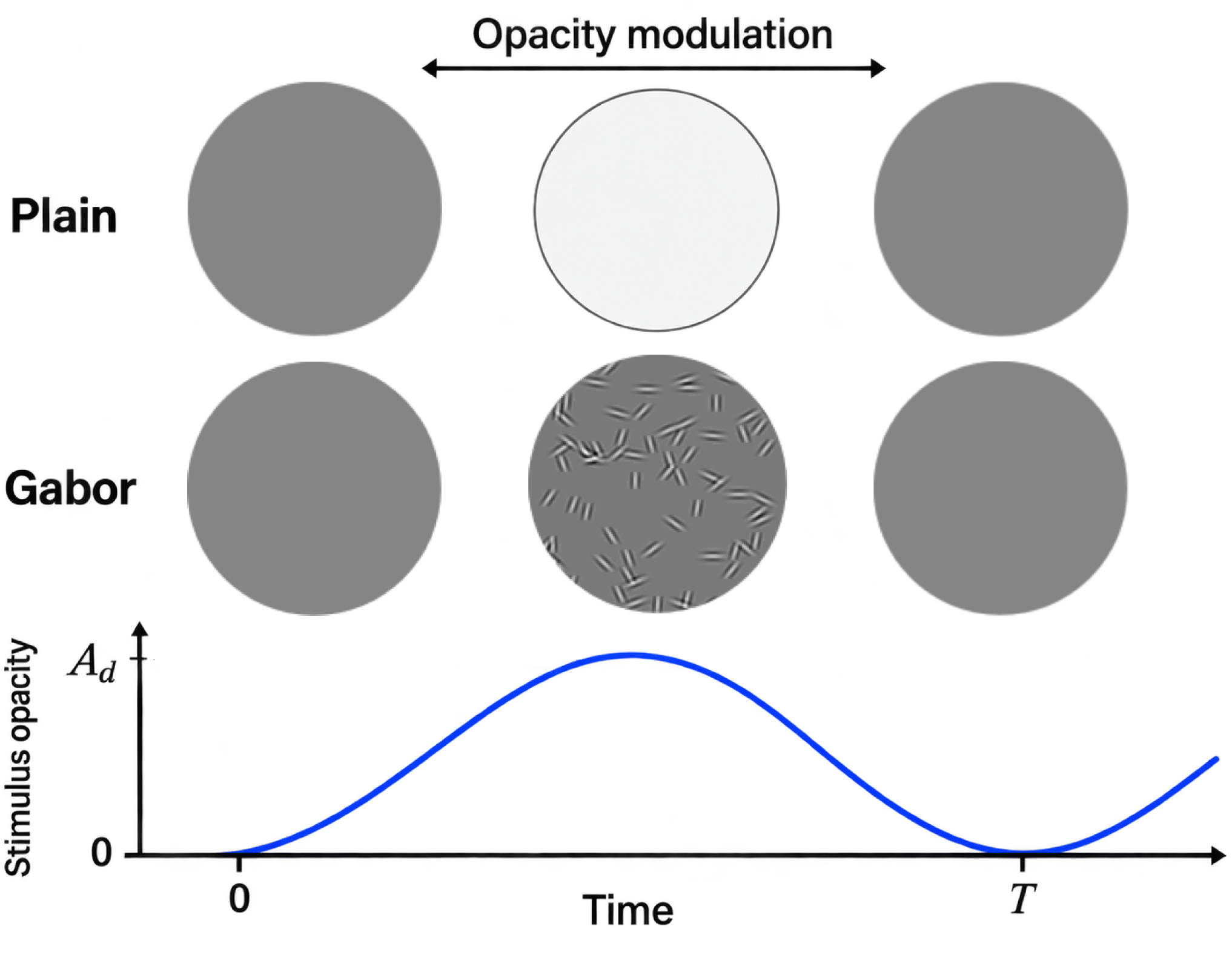
Types of flicker used in the experiment. Plain and Gabor stimuli were modulated continuously over time by varying stimulus opacity according to a sinusoidal function. The amplitude depth, *A_d_*, scaled the amplitude of this sinusoidal modulation and therefore controlled stimulus visibility. Stimulation duration and the total number of displayed frames were held constant across frequencies, while stimulation frequency determined the number of sinusoidal modulation cycles occurring within this fixed presentation period. The duration of one modulation cycle, or period *T*, was defined as *T* = 1*/f*, where *f* is the stimulation frequency in Hz. Adapted from Gomel et al. (2026).

Stimuli were generated in Python (version 3.8) and presented on a 24-inch LCD monitor (1920 *×* 1080 pixels, 60 Hz refresh rate). Stimuli consisted of 150 *×* 150 pixel circular disc presented at fixation, either as a uniform Plain stimulus or as a spatially textured Gabor stimulus. Participants sat approximately 1 meter from the screen and were instructed to maintain fixation while passively viewing the stimulation. The session began with a resting-state EEG recording used to estimate IAF. A perceptual threshold procedure was then conducted separately for Plain and Gabor stimuli. The main experiment comprised 12 blocks of 10 trials, with each trial consisting of 3 s of flicker followed by 3 s of rest. Visual comfort and mental fatigue were rated after each block. The session lasted approximately 50 min.

##### Staircase procedure: perceptual threshold

Perceptual visibility thresholds were estimated separately for Plain and Gabor stimuli using a two-phase staircase procedure. Participants viewed randomized blocks of four 1-s trials (two flickering and two static stimuli; 150 *×* 150 pixels) and reported whether flicker was detected. Amplitude depth first decreased from 20% in 2% steps until flicker could no longer be reliably identified, then increased in 1% steps until both flickering trials within a block were correctly detected. The resulting amplitude depth, *A_d_*, was taken as the perceptual visibility threshold.

#### 2.1.3 EEG acquisition and preprocessing

EEG data were streamed and temporally synchronized with behavioral events via Lab Streaming Layer (LSL) (Kothe et al., 2025). EEG data were recorded at 512 Hz using a 32-channel BioSemi ActiveTwo system with electrodes placed according to the international 10-20 system. Data preprocessing was conducted with EEGLAB (version 2024) and custom MATLAB (version R2024a) scripts. Data were re-referenced to the average reference and band-pass filtered between 1 and 20 Hz using a zero-phase finite impulse response (FIR) filter. Channels in which noise exceeded three standard deviations above the median power spectrum were detected *via* spectral thresholding, removed, and then interpolated by spherical spline interpolation from neighboring electrodes (mean = 0.79, SD = 0.73). Independent Component Analysis (ICA) was performed using the *Picard* algorithm to separate neural and non neural sources. Components reflecting non-neural artifacts (e.g., eye movements, muscle activity, cardiac signals, line noise or poor electrode contact) were automatically flagged by *ICLabel* (v1.5) and any component with *>* 70% confidence in an artifact category was excluded. On average, approximately 6.25 components (SD = 3.62) were removed per participant. Finally, the continuous EEG signal was segmented into epochs based on stimulus presentation. Each epoch spanned from *−*1000 to 4000 ms relative to the stimulus onset to prevent potential edge artifacts from contaminating the analysis window.

##### SSVEP: RESS Method

Rhythmic Entrainment Source Separation (RESS) was used to isolate stimulation-locked neural activity (Cohen & Gulbinaite, 2017). RESS derives a spatial filter by contrasting covariance matrices computed from EEG activity filtered around the stimulation frequency with covariance matrices obtained from adjacent frequencies. In our implementation, zero-phase FIR bandpass filters were applied around the individual target frequency and on flanking bands at *±* 1 Hz. Generalized eigenvalue decomposition was applied, and the eigenvector associated with the largest eigenvalue was selected as the RESS spatial filter. This filter was applied to the EEG data to obtain a single component per condition. SSVEP signal-to-noise ratio (SNR) was then quantified as the power at the stimulation frequency relative to the mean power in neighboring frequency bins.

##### Individual alpha frequency (IAF)

IAF was estimated from a 30-s closed-eyes resting-state EEG period using the automated approach of Corcoran et al. (2018). Data were band-pass filtered between 1 and 40 Hz and decomposed using ICA with the *Picard* algorithm. Given that rhythmic stimulation has the strongest effects in primary sensory cortices, particularly in response to low-level visual input such as flickering stimuli, we focused on the occipital alpha rhythm (Di Russo et al., 2007). Occipital components were then averaged into a single time series and spectral analysis was conducted throughout the 1–40 Hz range, with particular focus on the alpha band (8–13 Hz). A Savitzky-Golay filter was applied to smooth the spectrum, and power spectral density was estimated using the *pwelch* function. The *restingIAF* MATLAB function from (Corcoran et al., 2018) was subsequently used to determine the individual alpha peak.

#### 2.1.4 Questionnaires

At the end of each block, participants completed a series of questions designed to assess their subjective experience of the flickering stimuli. Specifically, the questionnaires evaluated visual comfort and mental fatigue. Responses were collected using an 11-point Likert scale ranging from 0 to 10. Visual comfort was rated from 0 (highly uncomfortable) to 10 (very comfortable), and mental fatigue from 0 (very fatigued) to 10 (not fatigued).

#### 2.1.5 Statistical analyses

Statistical analyses were performed in JASP (version 0.19.3). SSVEP SNR, visual comfort, and mental fatigue were analyzed using a 3 *×* 2 *×* 2 repeated-measures ANOVA with stimulation frequency (IAF *−*2 Hz *vs.* IAF *vs.* IAF +2 Hz), intensity (Full *vs.* Threshold), and flicker type (Plain *vs.* Gabor) as within-subject factors. Normality was assessed using the Shapiro–Wilk test, sphericity was assessed using Mauchly’s test, and pairwise post hoc comparisons were corrected for multiple comparisons using Holm’s procedure (Holm, 1979).

### 2.2 Results

#### 2.2.1 SSVEP signal quality

SSVEP signal quality was strongly influenced by stimulation frequency (*F* (2, 66) = 24.942, *p <* 0.001, *η*^2^ = 0.126). SNR was highest at the individual alpha frequency (IAF; *M* = 7.23, *SD* = 2.84), with significantly higher responses than at IAF*−*2 Hz (*M* = 4.52, *SD* = 1.85; *p <* 0.001, *d* = 0.775) and IAF+2 Hz (*M* = 4.81, *SD* = 1.79; *p <* 0.001, *d* = 0.693). Full-intensity stimulation (*M* = 6.32, *SD* = 2.96) also produced higher SNR than threshold-intensity stimulation (*M* = 4.72, *SD* = 1.30), *F* (1, 33) = 9.303, *p* = 0.004, *η*^2^ = 0.055).

There was no overall main effect of flicker type (*F* (1, 33) = 0.038, *p* = 0.856), indicating comparable SNR for Plain (*M* = 5.49, *SD* = 1.88) and Gabor stimulation (*M* = 5.55, *SD* = 1.86). A significant Frequency *×* Flicker Type interaction (*F* (2, 66) = 8.110, *p <* 0.001, *η*^2^ = 0.032), indicated that frequency detuning affected the two stimulus formats differently. Post-hoc comparisons showed that Gabor flicker exhibited higher SNR at IAF (*M* = 7.58, *SD* = 3.83) than at both IAF*−*2 (*M* = 5.08, *SD* = 2.68; *p* = 0.008, *d* = 0.715) and IAF+2 (*M* = 3.98, *SD* = 1.53; *p <* 0.001, *d* = 1.031). For Plain flicker, SNR was higher at IAF (*M* = 6.88, *SD* = 2.69) than at IAF*−*2 (*M* = 3.97, *SD* = 1.52; *p <* 0.001, *d* = 0.835), whereas the IAF versus IAF+2 (*M* = 5.64, *SD* = 2.94) comparison did not reach significance (*p* = 0.054, *d* = 0.355).

Finally, a significant Frequency *×* Flicker Type *×* Intensity interaction was also observed (*F* (2, 66) = 5.907, *p* = 0.004, *η*^2^ = 0.016). Post-hoc analyses at the IAF revealed that Gabor and Plain flickers produced comparable SNR at both Full intensity (Gabor: *M* = 8.55, *SD* = 5.32; Plain: *M* = 7.56, *SD* = 3.94; *p >* 0.05) and Threshold intensity (Gabor: *M* = 6.60, *SD* = 4.09; Plain: *M* = 6.19, *SD* = 2.91; *p >* 0.05), indicating that Gabor stimulation did not compromise neural responsiveness when delivered at the individual alpha frequency. The interaction effects mainly reflected differences in the sensitivity of the two stimulus formats to frequency detuning. For Gabor flicker, SNR was lower at IAF +2 (*M* = 4.04, *SD* = 1.34) Hz than at IAF under full intensity (*M* = 8.55, *SD* = 5.32; *p* = 0.001, *d* = 1.293). For Plain flicker, SNR was lower at IAF*−*2 than at IAF under both Full intensity (IAF*−*2: *M* = 4.43, *SD* = 2.43; IAF: *M* = 7.56, *SD* = 3.94; *p <* 0.001, *d* = 0.897) and Threshold intensity (IAF*−*2: *M* = 3.50, *SD* = 1.29; IAF: *M* = 6.19, *SD* = 2.91; *p* = 0.001, *d* = 0.774; Figure 2).

**Figure 2:**
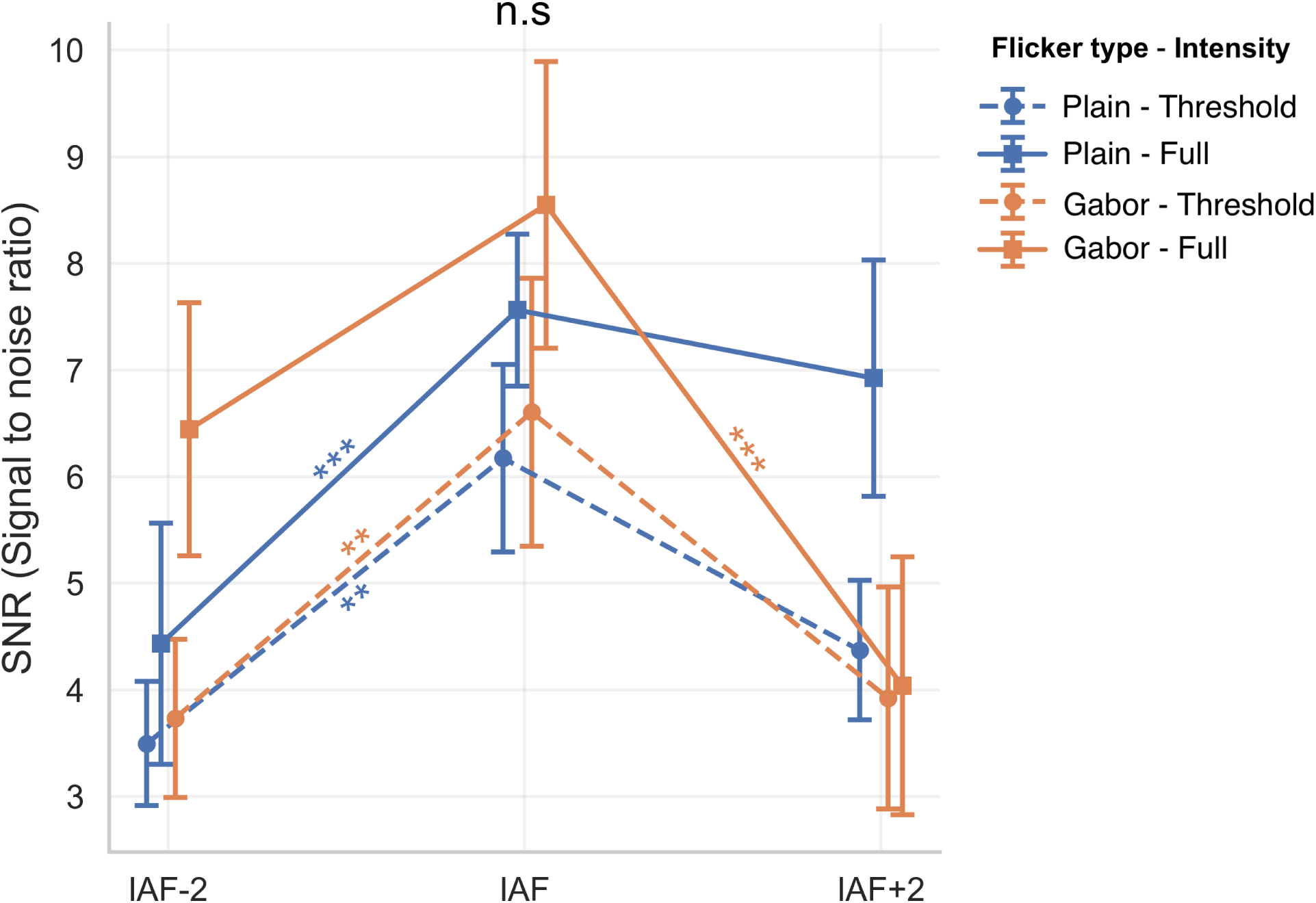
SSVEP signal-to-noise ratio (SNR) as a function of flicker type, stimulation frequency, and intensity. Vertical bars represent 95% confidence intervals. Asterisks indicate significant pairwise comparisons between IAF and neighboring frequencies within a given flicker type and intensity condition (*** *p <* 0.001; ** *p <* 0.01; n.s., non-significant.)

#### 2.2.2 Subjective measures Visual comfort

Subjective ratings indicated that visual comfort was strongly influenced by stimulation parameters, with a clear advantage for Gabor flicker (see Supplementary Figure S1). A significant main effect of Flicker Type (*F* (1, 33) = 81.602, *p <* 0.001, *η*^2^ = 0.260) showed that Gabor stimulation (*M* = 6.77, *SD* = 1.16) was rated as more comfortable than Plain stimulation (*M* = 4.34, *SD* = 1.32). Visual comfort was also affected by Frequency (*F* (2, 66) = 13.756, *p <* 0.001, *η*^2^ = 0.021), with the highest comfort ratings at IAF (*M* = 5.99, *SD* = 1.10), compared with IAF*−*2 (*M* = 5.15, *SD* = 1.16) and IAF+2 (*M* = 5.52, *SD* = 1.05). A significant main effect of Intensity was also observed (*F* (1, 33) = 41.664, *p <* 0.001, *η*^2^ = 0.136), with higher comfort ratings at Threshold intensity (*M* = 6.43, *SD* = 1.29) than at Full intensity (*M* = 4.67, *SD* = 1.20).

These main effects were qualified by significant Flicker Type *×* Intensity (*F* (1, 33) = 42.556, *p <* 0.001, *η*^2^ = 0.039), and Flicker Type *×* Frequency interactions (*F* (2, 66) = 21.095, *p <* 0.001, *η*^2^ = 0.049). A significant Flicker Type *×* Intensity *×* Frequency interaction was also observed (*F* (2, 66) = 7.325, *p* = 0.001, *η*^2^ = 0.015). Post-hoc analyses revealed that at IAF, Gabor flicker was rated as more comfortable than Plain flicker at both Full intensity (Gabor: *M* = 7.82, *SD* = 1.68; Plain: *M* = 2.29, *SD* = 1.59; *p <* 0.001, *d* = *−*2.910) and Threshold intensity (Gabor: *M* = 7.97, *SD* = 1.91; Plain: *M* = 5.85, *SD* = 1.81; *p <* 0.001, *d* = *−*1.151). Reducing intensity markedly improved comfort for Plain flicker (Full: *M* = 2.29, *SD* = 1.59; Threshold: *M* = 5.85, *SD* = 1.81; *p <* 0.001, *d* = *−*1.839), whereas comfort ratings remained similar between Full (*M* = 7.82, *SD* = 1.68) and Threshold intensity (*M* = 7.97, *SD* = 1.91) for Gabor flicker (*p* = 1.000).

##### Mental Fatigue

Mental fatigue showed a comparable pattern. A significant main effect of Flicker Type (*F* (1, 33) = 18.313, *p <* 0.001, *η*^2^ = 0.067), indicated higher ratings for Gabor (*M* = 6.20, *SD* = 1.78) than Plain flicker (*M* = 5.16, *SD* = 1.66), reflecting lower subjective fatigue with Gabor stimulation. Mental fatigue ratings were also affected by Frequency (*F* (2, 66) = 17.163, *p <* 0.001, *η*^2^ = 0.037), with ratings highest at IAF (*M* = 6.10, *SD* = 1.56), followed by IAF+2 (*M* = 5.78, *SD* = 1.75) and IAF*−*2 (*M* = 5.16, *SD* = 1.66). A significant main effect of Intensity was also observed (*F* (1, 33) = 9.249, *p* = 0.005, *η*^2^ = 0.029), with higher ratings at Threshold intensity (*M* = 6.03, *SD* = 1.78) than at Full intensity (*M* = 5.33, *SD* = 1.62). Two-way and three-way interactions also reached significance.

Overall, Gabor flicker was rated as less mentally fatiguing than Plain flicker (see Supplementary Figure S2). Altogether, these results indicate that visual comfort and mental fatigue are jointly modulated by spatial pattern, flicker intensity, and temporal frequency, with Gabor patterns producing the most comfortable and least fatiguing perceptual experience, particularly at the IAF, making it particularly suitable for subsequent cognitive and neuromodulation paradigms requiring sustained rhythmic stimulation.

#### 2.2.3 Integrated view of neural and subjective measures

To provide an integrated view of these findings, Figure 3 summarizes the joint relationship between SNR, visual comfort, and mental fatigue across experimental conditions. Gabor flicker at IAF occupied the region associated with the most favorable overall balance: relatively high SNR, high comfort, and low fatigue. Although Plain flicker at Full intensity could produce comparable SNR values, it was associated with poorer subjective ratings. This integrative pattern suggests that individualized Gabor flicker provides the most suitable compromise between entrainment efficacy and tolerability.

**Figure 3:**
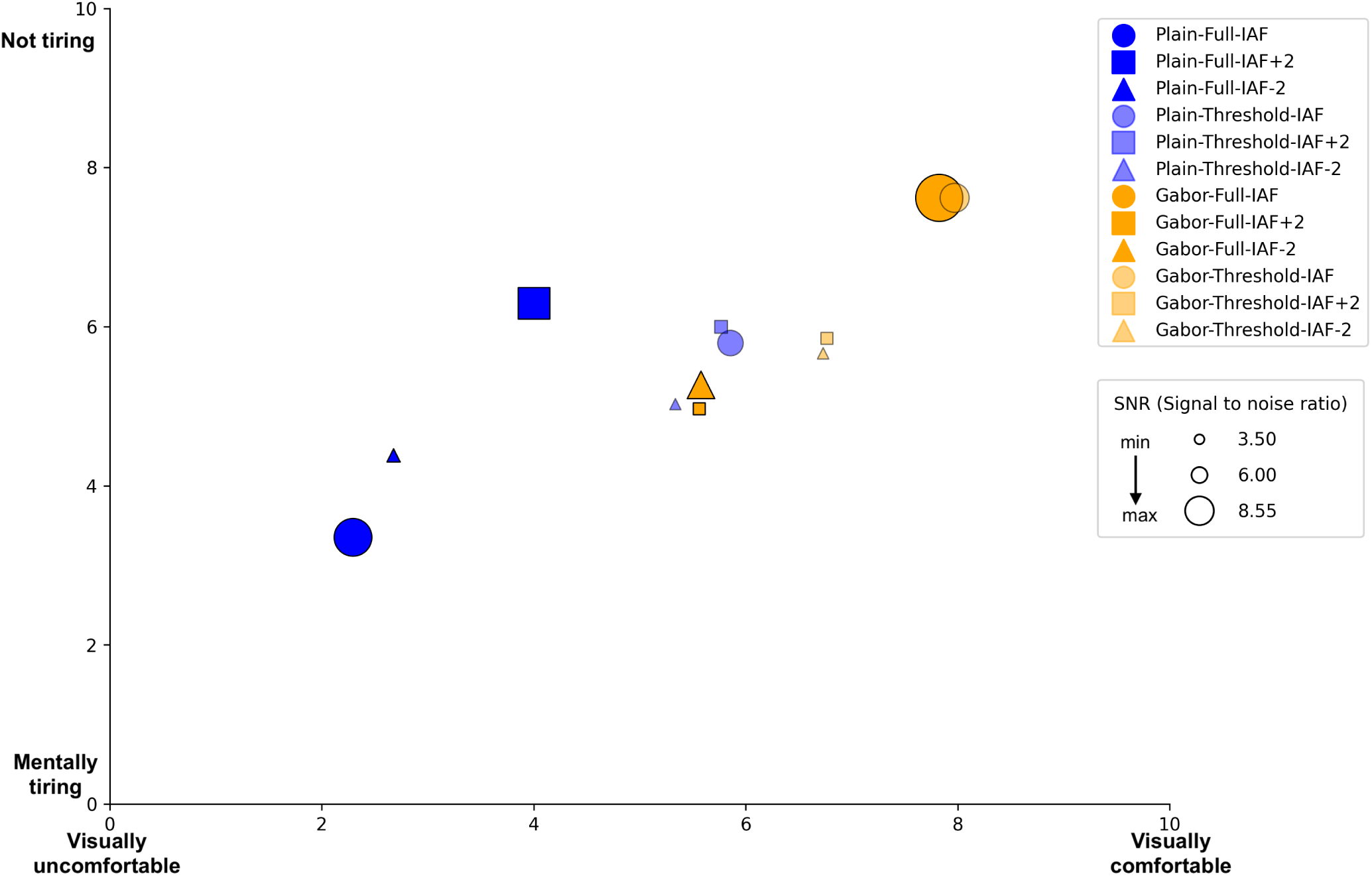
Relationship between subjective visual comfort (x axis), mental fatigue (y axis), and SNR across experimental conditions. Point color denotes Flicker Type (Plain vs. Gabor), marker shape denotes stimulation Frequency (IAF - 2 Hz, IAF, IAF + 2 Hz), and transparency indicates Intensity (Full vs. Threshold). Marker size is proportional to mean SNR. Higher values indicate better comfort and lower fatigue.

### 2.3 Discussion

Experiment 1 examined how frequency, intensity, and stimulus structure jointly shaped visual entrainment and subjective tolerability. The main finding was that SSVEP signal quality was maximal when stimulation was delivered at each participant’s individual alpha frequency, whereas small deviations from IAF reduced SNR. This finding is consistent with the view that rhythmic visual stimulation is more effective when the driving frequency is aligned with endogenous oscillatory dynamics (Gulbinaite et al., 2017; Keitel et al., 2019; Notbohm & Herrmann, 2016). In other words, stimulation at the IAF may interact more efficiently with ongoing alpha dynamics because the external input is temporally aligned with the natural frequency of the underlying neural system. Rather than imposing an entirely new rhythm, the flicker may therefore reinforce or amplify an already preferred oscillatory mode, leading to stronger frequency-specific neural responses. It also supports the relevance of individualized stimulation protocols for visual neuromodulation, as fixed frequency stimulation may fail to optimally engage alpha-band dynamics in all participants (De Graaf & Duecker, 2022; Souza & Oberauer, 2016).

Importantly, neural efficacy was not determined by frequency alone. Intensity and stimulus structure also contributed to the balance between entrainment strength and subjective experience. Full intensity stimulation generally increased SNR, but threshold-adjusted stimulation was better tolerated. Moreover,

Gabor and Plain flickers produced comparable SNR at the IAF, while Gabor stimulation was consistently rated as more comfortable and less mentally fatiguing. Notably, the comfort penalty associated with full-intensity stimulation was primarily observed for Plain flicker, suggesting that Gabor stimulation can preserve strong frequency-specific neural responses while reducing the perceptual burden associated with conventional flicker (Dehais et al., 2024; Ladouce & Dehais, 2024).

Overall, this first experiment identified individualized Gabor flicker at IAF as the most favorable compromise between neural efficacy and subjective tolerability. These findings provide the methodological basis for Experiment 2, in which the same individualized and visually comfortable stimulation approach was extended to alpha- and theta-frequency stimulation during working-memory retention.

## 3 EXPERIMENT 2

### 3.1 Materials and method

#### 3.1.1 Participants

Thirty four healthy adults participated in experiment 2 (11 women; mean age = 23 years, SD = 3.80 years). Only three participants had also taken part in Experiment 1. All reported normal or corrected-to-normal visual acuity and no history of neurological or psychiatric disorders. Participants provided written informed consent; the protocol was approved by the local ethics committee (CER No. 2025-1205) in accordance with the Declaration of Helsinki. Two participants were excluded from the analyses: one due to EEG recording issues, characterized by poor signal quality likely caused by electrode disconnection, and one because they did not complete the experiment. The final sample therefore included thirty two participants (11 women; mean age = 23.15 years, SD = 3.86 years). Participants were compensated €10 for participating in the study.

#### 3.1.2 Stimuli and Experimental Paradigm

The experimental paradigm was adapted from Riddle et al. (2020) and consisted of a modified delayed match-to-sample task with a retrospective spatial cue (retro-cue; Figure 4). An example of the experimental procedure is shown in Supplementary Video S1. The task was programmed in Python (version 3.9). Stimuli were generated in Python (version 3.9) and presented on a 24-inch LCD monitor (1920 *×* 1080 pixels, 60 Hz) at a viewing distance of approximately 1 m. Based on Experiment 1, rhythmic stim-ulation was delivered using Gabor-textured (Figure 1) overlays covering the two memory regions. Before the task, participants completed a 5-min resting-state EEG recording used to estimate individual alpha and theta stimulation frequencies. They then completed 48 training trials followed by six experimental blocks of 56 trials.

**Figure 4:**
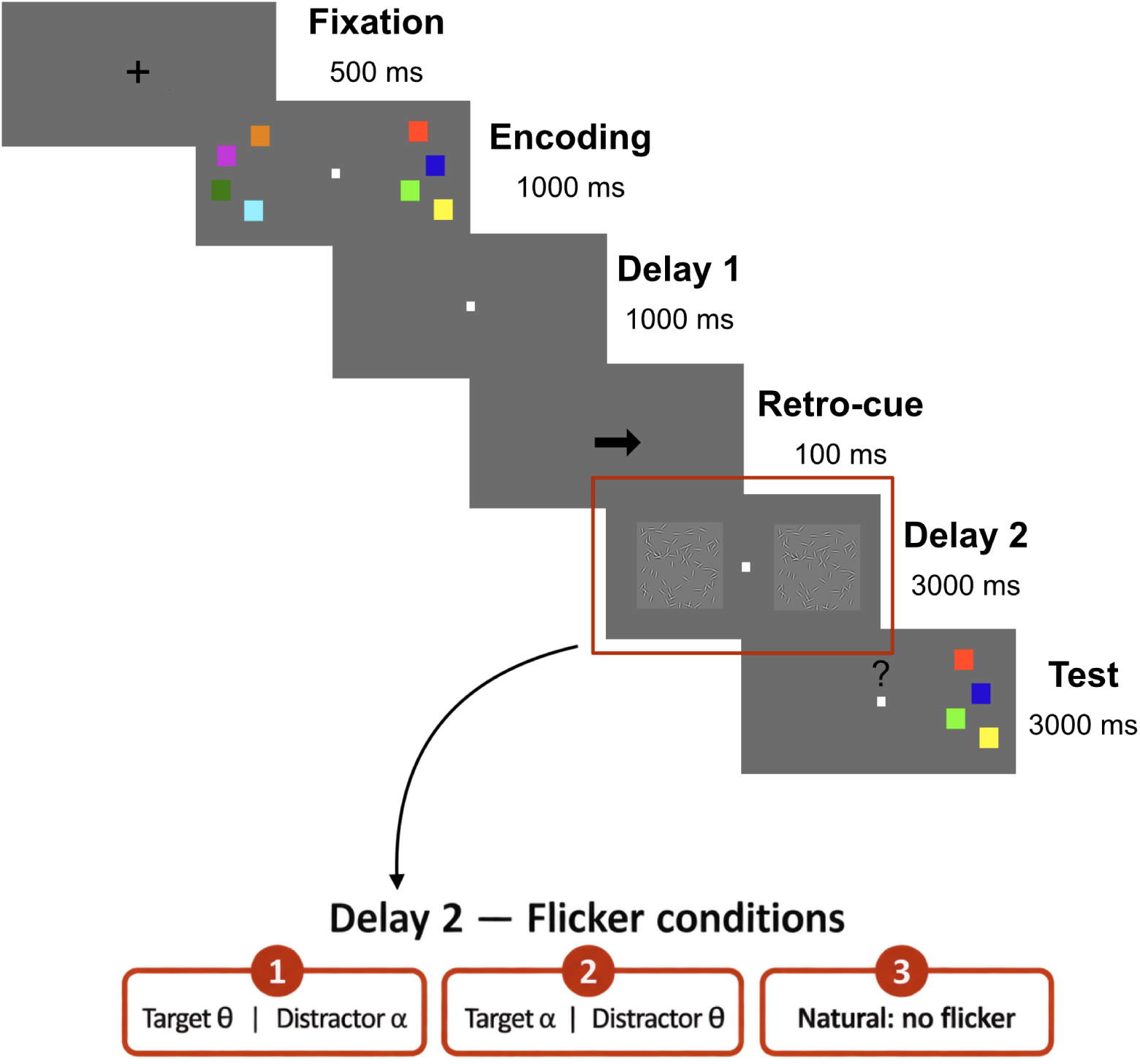
Experimental procedure. Schematic illustration of a trial in the memory load 4 condition. The experiment included two memory loads (4 and 5 items per hemifield) and three stimulation conditions during Delay 2 (*T_theta_*/*D_alpha_*), (*T_alpha_*/*D_theta_*), and Natural (no flicker). Target refers to the hemifield indicated by the retro-cue arrow (pointing left or right), whereas Distractor refers to the opposite, nontarget hemifield.

Each block implemented one of three stimulation conditions: (1) a Target-theta and Distractor-alpha condition (*T_theta_*/*D_alpha_*), in which individual theta-frequency flicker (ITF) was applied to the cued hemifield and alpha-frequency flicker (IAF) to the uncued hemifield (see section 3.1.3 for IAF and ITF evaluations); (2) a Target-alpha and Distractor-theta condition (*T_alpha_*/*D_theta_*), corresponding to the reverse frequency arrangement; and (3) a Natural condition, in which Gabor stimuli were presented without rhythmic flicker. Flicker onset was time locked to retro-cue offset and continued throughout the second retention interval. On neutral-cue trials, both stimulus regions flickered at 30 Hz.

Each trial began with the presentation of a central fixation cross for 500 ms, followed by a bilateral encoding array displayed for 1000 ms. The encoding array consisted of colored squares presented simultaneously in two rectangular regions located symmetrically to the left and right of fixation and each item was equidistant from fixation. Following a first retention delay of 1000 ms, either a retro-cue (50% of trials) or a neutral cue (50% of trials) was presented centrally for 100 ms. Retro-cues were 100% predictive of the hemifield that would be tested at the end of the trial and were presented as leftward or rightward arrows indicating which hemifield should be prioritized in memory. Neutral cues took the form of a bilateral double arrow, indicating that both hemifields should be maintained with equal priority. Following the cue, a second retention delay of 3000 ms was presented, during which textured flicker was applied according to the block condition. At the end of this delay, participants were probed with a test array presented either to the left or to the right of fixation, at the same spatial location as the original encoding display 4. The test array was identical to the encoding array (match trial) or differed by a single color change affecting one item (non-match trial). On non-match trials, the replacement color was selected to maximize discriminability from the original colors and was drawn from the set of colors not present in the tested memory array. Participants were instructed to respond as accurately and as quickly as possible by pressing the “K” keyboard key for same or “D” for different within a 3000 ms response window.

Stimuli were displayed on a uniform gray background (RGB : 126, 126, 126), and the stimulus set comprised nine colors. Depending on the memory load, four or five items were presented in each hemifield. Items were pseudo-randomly positioned within each rectangular region under spacing constraints ensuring at least one empty-item gap between neighboring positions. Memory load and cue type (retro vs. neutral) were intermixed, randomized, and balanced across trials.

#### 3.1.3 EEG Acquisition and preprocessing

Continuous EEG data were recorded at 500 Hz using a 32-channel MBT semi-dry EEG system (saline soaked sponges - MbrainTrain) with electrodes positioned according to the international 10–20 system. EEG data were streamed and synchronized with behavioral events via LSL (Kothe et al., 2025). Data preprocessing followed the same procedure as in Experiment 1. Regarding ICA decomposition, on average, 5.41 components (*SD* = 3.42) were excluded per participant. Data were then epoched from *−*1000 to 4000 ms relative to the onset of the second retention interval (delay2).

##### Individualized Rhythmic Visual Stimulation

Stimulation frequencies were individualized from the resting-state EEG recorded before the task. IAF was estimated using the procedure described in Experiment 1 (see section 2.1.3). Individual theta frequency (ITF) was estimated from a fronto-central region of interest comprising Fz, FC1, FC2, and Cz (Smit et al., 2023). Power spectral density was computed using Welch’s method with a 4-s Hamming window, 50% overlap, and zero-padding to the next power of two. ITF was defined as the frequency showing maximal power within the 4-8 Hz range.

##### SSVEP Analysis and Signal to Noise Ratio

SSVEP signal-to-noise ratio (SNR) was computed following the same RESS pipeline as in Experiment 1. Spatial filters were estimated from frequency-specific and flanking-band covariance matrices and then applied to the EEG data to extract a single component per condition. SNR was subsequently quantified for each participant in each of the three block types (*T_theta_*/*D_alpha_*, *T_alpha_*/*D_theta_* and Natural), allowing direct comparison of rhythmic entrainment across stimulation conditions.

To limit contamination by unusually strong responses at the distractor frequency, an additional trial-level rejection procedure was applied. For each participant, RESS filters derived when a given frequency was presented on the target side were applied to trials in which that same frequency was presented on the distractor side. Distractor-frequency SNR was then computed trial by trial. An individualized rejection threshold was defined from the 56 retro-cue trials of the Natural condition as the mean SNR plus three standard deviations. This threshold was then applied to the 112 retro-cue trials from the four rhythmic-stimulation blocks (28 retro-cue trials per block, with two memory-load levels for each stimulation condition). Trials whose distractor frequency SNR exceeded this threshold were excluded from EEG analyses, together with the corresponding behavioral trials. On average, 6.50 of these 112 trials per participant were excluded (*SD* = 6.65).

##### Inter-trial phase coherence

To further characterize the temporal consistency of rhythmic entrainment during working memory retention, inter-trial phase clustering (ITPC) was computed during the second retention interval on retro-cue trials only. ITPC provides a measure of trial-to-trial phase consistency and is commonly used to quantify the extent to which neural activity aligns in phase across repeated events (Bastos & Schoffelen, 2016). Time-frequency analyses were performed in Python using MNE-Python and complex Morlet wavelets (tfr array morlet). Following standard phase-based time-frequency approaches (Cohen, 2014), complex coefficients were normalized to unit magnitude and averaged across trials, with ITPC defined as the magnitude of the resulting mean phase vector. Analyses were restricted to occipital channels (O1 and O2), as visually evoked steady-state responses are typically strongest over posterior-occipital sites (Norcia et al., 2015). ITPC was computed separately for each stimulation condition and memory load and compared with the Natural condition.

#### 3.1.4 Behavioral responses

Participants’ accuracy and response speed were recorded. For each participant, reaction times (RTs) were extracted on a trial-by-trial basis. Trials were excluded if they met any of the following criteria: no response or RTs longer than 3000 ms, or RTs shorter than 200 ms. The remaining RTs were then averaged for each participant across block stimulation (*T_theta_*/*D_alpha_ vs. T_alpha_*/*D_theta_ vs.* Natural), and memory load (4 items *vs.* 5 items).

On average, 140 of the 168 retro-cue trials per participant (28 retro-cue trials for each stimulation condition and memory load; *SD* = 20.01) were retained after applying all exclusion criteria, including EEG-based rejection and behavioral filtering.

#### 3.1.5 Statistical analyses

Statistical analyses were performed in JASP (version 0.19.3). Accuracy and RT were first analyzed using a 3 *×* 2 *×* 2 repeated-measures ANOVA with stimulation condition (*T_theta_*/*D_alpha_*, *T_alpha_*/*D_theta_* and Natural), memory load (4 vs. 5 items), and cue type (retro vs. neutral) as within-subject factors. To specifically assess stimulation effects, analyses were then restricted to retro-cue trials. Retro-cues selectively prioritize one representation maintained in working memory while making the opposite representation task-irrelevant, thereby providing a well established framework for studying attentional prioritization within working memory (Riddle et al., 2020; Souza & Oberauer, 2016). The retro-cue analyses were therefore conducted using 3 *×* 2 repeated-measures ANOVAs with stimulation condition and memory load as factors. The same 3 *×* 2 design was applied to SNR and ITPC. Normality was assessed using the Shapiro–Wilk test, sphericity was assessed using Mauchly’s test, and significant effects were followed up with paired-samples *t*-tests using Holm correction for multiple comparisons (Holm, 1979).

### 3.2 Results

#### 3.2.1 SSVEPs analyses

Rhythmic stimulation produced robust frequency-specific responses during working-memory retention. SNR differed significantly across conditions (*F* (2, 62) = 30.003, *p <* 0.001, *η*^2^ = 0.440). SNR was higher during ITF stimulation (*M* = 17.08, *SD* = 9.12; *p <* 0.001, *d* = 1.496) and IAF stimulation (*M* = 9.42, *SD* = 7.53; *p <* 0.001, *d* = 0.782) than in the Natural condition (*M* = 1.02, *SD* = 0.17; Figure 5).

**Figure 5:**
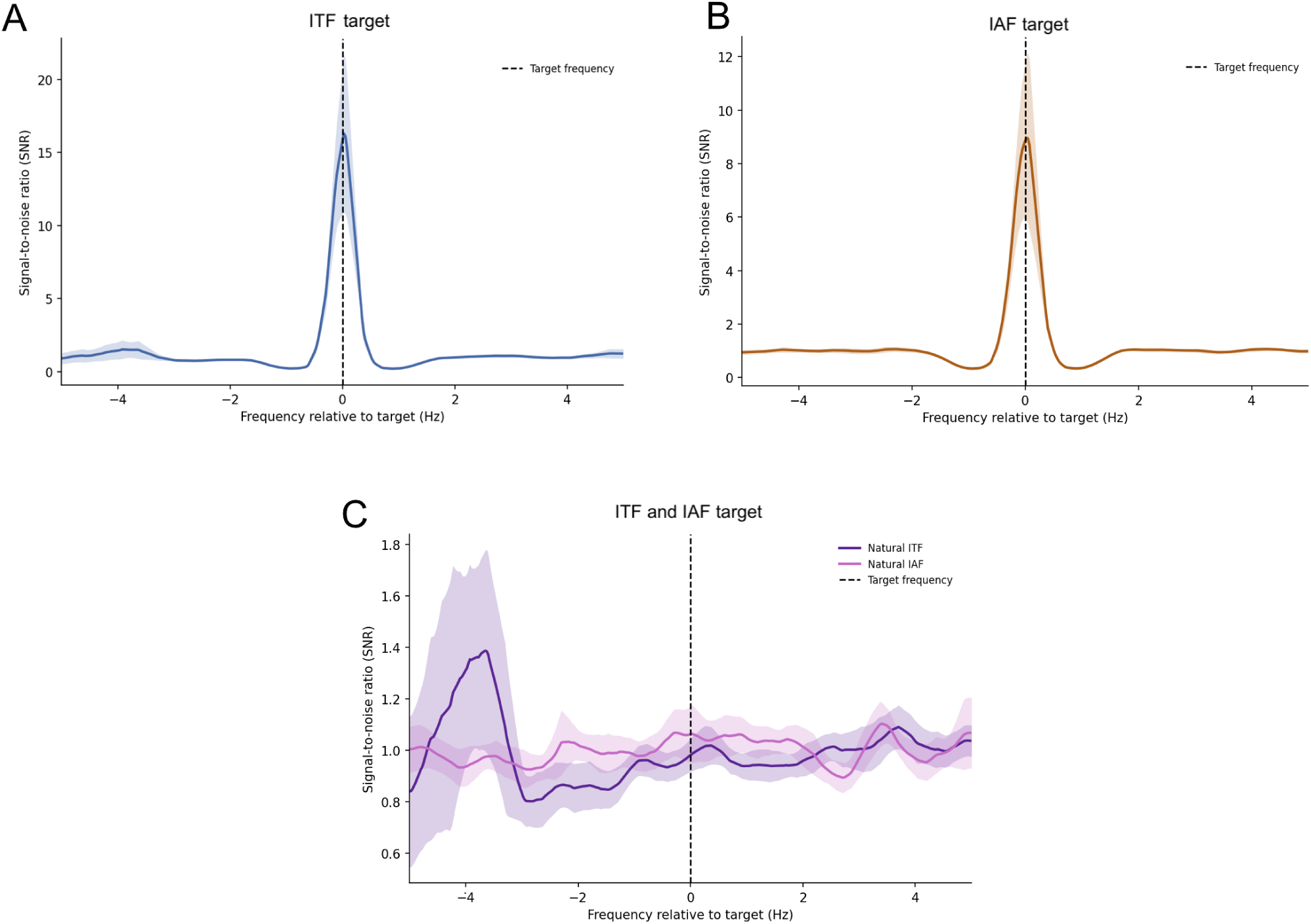
Mean signal-to-noise ratio (SNR) as a function of frequency relative to the individual target frequency for each stimulation condition. SNR values are plotted relative to the target stimulation frequency, which is centered at 0 on the x-axis; negative and positive values indicate frequencies below and above the target frequency, respectively. **A**. *T_theta_*/*D_alpha_* condition, in which the target hemifield flickered at the individual theta frequency (ITF) and the distractor hemifield at the individual alpha frequency (IAF). **B**. *T_alpha_*/*D_theta_* condition, in which the target hemifield flickered at IAF and the distractor hemifield at ITF. **C**. Natural condition, in which no rhythmic visual stimulation was applied and SNR therefore reflected baseline endogenous activity around the corresponding frequencies. Solid lines represent mean SNR values across participants, and shaded areas indicate the 95% confidence intervals. The pronounced peak at the frequency centered on 0 in the stimulation conditions reflects the frequency-tagged steady-state visual evoked response.

ITPC showed a similar pattern, with a significant effect of stimulation condition (*F* (2, 62) = 152.900, *p <* 0.001, *η*^2^ = 0.756, Figure 6). ITPC was higher during ITF (*T_theta_*/*D_alpha_*; *M* = 0.815, *SD* = 0.141) than IAF stimulation (*T_alpha_*/*D_theta_*; *M* = 0.659, *SD* = 0.157; *p <* 0.001, *d* = 1.090), and both rhythmic stimulation conditions produced greater ITPC than the Natural condition (both *p <* 0.001). Neither memory load nor the interaction between load and stimulation condition was significant.

**Figure 6:**
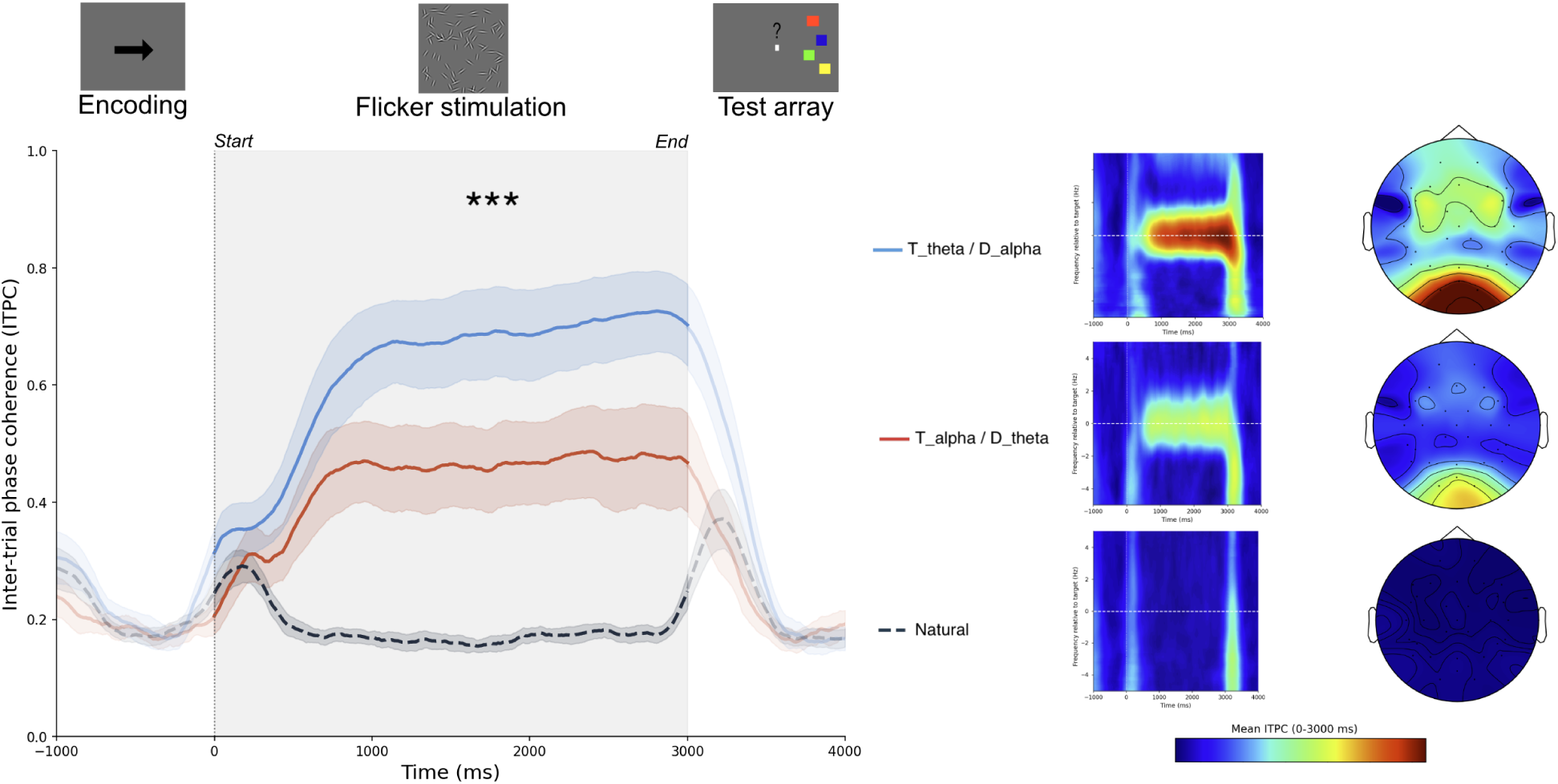
Inter-trial phase coherence (ITPC) dynamics across stimulation conditions. The left panel depicts the ITPC time course averaged across occipital electrodes (O1/O2) for the three experimental conditions (*T_theta_*/*D_alpha_*, *T_alpha_*/*D_theta_*, and Natural). Colored traces show the group mean, with shaded envelopes indicating the 95% confidence intervals. The right panels present the corresponding time-frequency representations and scalp topographies for each condition, highlighting the spectral and spatial distribution of phase coherence. In the time–frequency plots, the dashed horizontal line denotes 0 Hz relative to the individual target frequency. Scalp topographies were obtained by averaging ITPC within the analysis window (0 - 3000ms). Warmer colors correspond to greater phase consistency across trials.

Taken together, these electrophysiological findings show that individualized Gabor flicker reliably entrained neural activity during working memory retention, as indicated by increased SNR and ITPC relative to the non rhythmic Natural condition.

#### 3.2.2 Behavioral performance

We next examined whether these neural effects were accompanied by changes in behavioral performance, beginning with the effects of cue type on accuracy and reaction time. Accuracy was significantly affected by memory load (*F* (1, 31) = 9.014, *p* = 0.005, *η*^2^ = 0.040), with higher accuracy at Load 4 (*M* = 63.81%, *SD* = 7.32) than at Load 5 (*M* = 60.29%, *SD* = 5.41). In contrast, the main effect of cue type on accuracy was not significant (*F* (1, 31) = 2.074, *p* = 0.160, *η*^2^ = 0.005), although accuracy was numerically higher on retro-cue trials (*M* = 62.70%, *SD* = 5.92) than on neutral-cue trials (*M* = 61.39%, *SD* = 6.30). For reaction times, only the main effect of cue type reached significance (*F* (1, 31) = 38.379, *p <* 0.001, *η*^2^ = 0.092), with faster responses following retro-cues (*M* = 1.206 s, *SD* = 0.264) than neutral cues (*M* = 1.309 s, *SD* = 0.280) (see Supplementary Figure S3). The main effect of memory load approached significance (*F* (1, 31) = 3.947, *p* = 0.056, *η*^2^ = 0.023), with numerically faster responses at Load 4 (*M* = 1.232 s, *SD* = 0.265) than at Load 5 (*M* = 1.283 s, *SD* = 0.289).

To evaluate the effect of rhythmic visual stimulation on behavioral performance, subsequent analyses were restricted to retro-cue trials, as these were the only trials in which individualized alpha or theta frequency flicker was applied to the target or distractor depending on the stimulation block. As shown in Figure 7, accuracy was significantly modulated by memory load (*F* (1, 31) = 4.923, *p* = 0.034, *η*^2^ = 0.039), with participants showing higher accuracy at Load 4 (*M* = 64.53%, *SD* = 8.29) than at Load 5 (*M* = 60.87%, *SD* = 6.71). In contrast, neither the main effect of stimulation block (*F* (2, 62) = 0.060, *p* = 0.941), nor the interaction between stimulation block and memory load (*F* (2, 62) = 0.036, *p* = 0.965), reached significance. Accuracy was descriptively similar across the Natural (*M* = 62.35%, *SD* = 7.21), *T_theta_*/*D_alpha_* (*M* = 62.78%, *SD* = 8.78), and *T_alpha_*/*D_theta_* (*M* = 62.97%, *SD* = 9.10) conditions (see Supplementary Figure S4). Similarly, no significant main effects or interactions were observed for reaction times, which were comparable across Natural (*M* = 1.182 s, *SD* = 0.275), T*_θ_*/D*_α_* (*M* = 1.217 s, *SD* = 0.290), and *T_alpha_*/*D_theta_* (*M* = 1.221 s, *SD* = 0.269) conditions (see Supplementary Figure S5).

**Figure 7:**
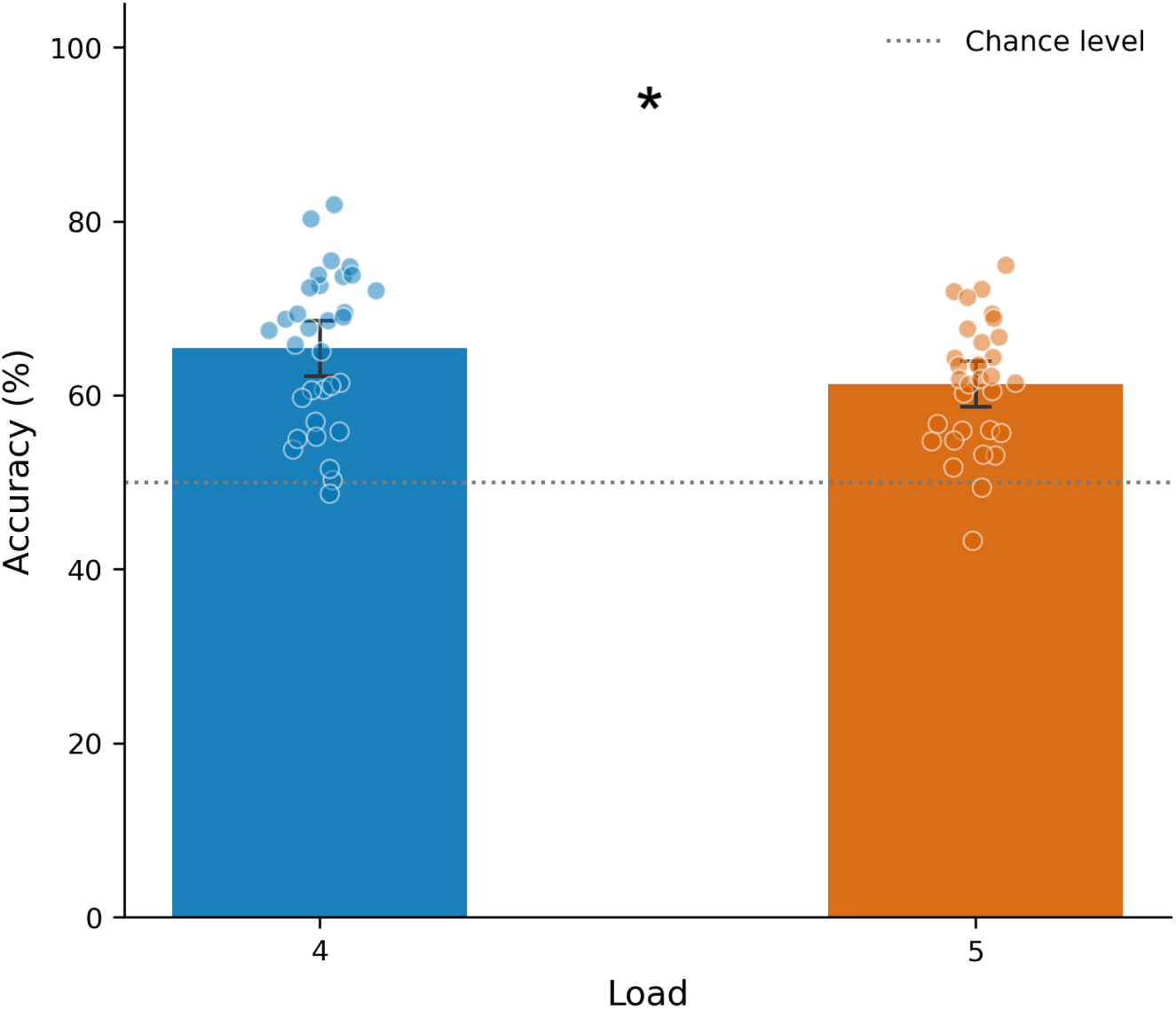
Accuracy on retro-cue trials as a function of memory load. Each point represents an individual participant value. Error bars indicate 95% confidence intervals. Asterisks indicate significant differences (* *p <* 0.05).

##### Exploratory analysis of baseline cue benefit

Given the substantial inter-individual variability in behavioral performance, we further examined whether stimulation effects depended on baseline cognitive efficiency. Previous studies have shown that the behavioral effects of rhythmic or sensory stimulation can be modest, context dependent, and more pronounced in participants with lower baseline working-memory performance or capacity (Pileckyte & Soto-Faraco, 2024; Riddle et al., 2020; Souza & Oberauer, 2016). We therefore conducted an exploratory post hoc analysis in participants who did not show a retro-cue benefit in the Natural condition at load 4 (*N* = 15), reasoning that these participants may have had greater scope for stimulation-related improvement because endogenous cue-based prioritization was less effective.

In this subgroup, a 3 *×* 2 repeated-measures ANOVA revealed a significant main effect of stimulation condition on accuracy (*F* (2, 28) = 3.631, *p* = 0.040, *η*^2^ = 0.075). Post hoc comparisons showed higher accuracy in the *T_theta_*/*D_alpha_* condition (*M* = 65.07%, *SD* = 7.44) than in the Natural condition (*M* = 60.02%, *SD* = 6.35; *p* = 0.040, *d* = 0.499), whereas the comparison between *T_theta_*/*D_alpha_* and *T_alpha_*/*D_theta_* (*M* = 60.10%, *SD* = 8.25) did not reach significance (*p* = 0.084*, d* = 0.492; Figure 8).

**Figure 8:**
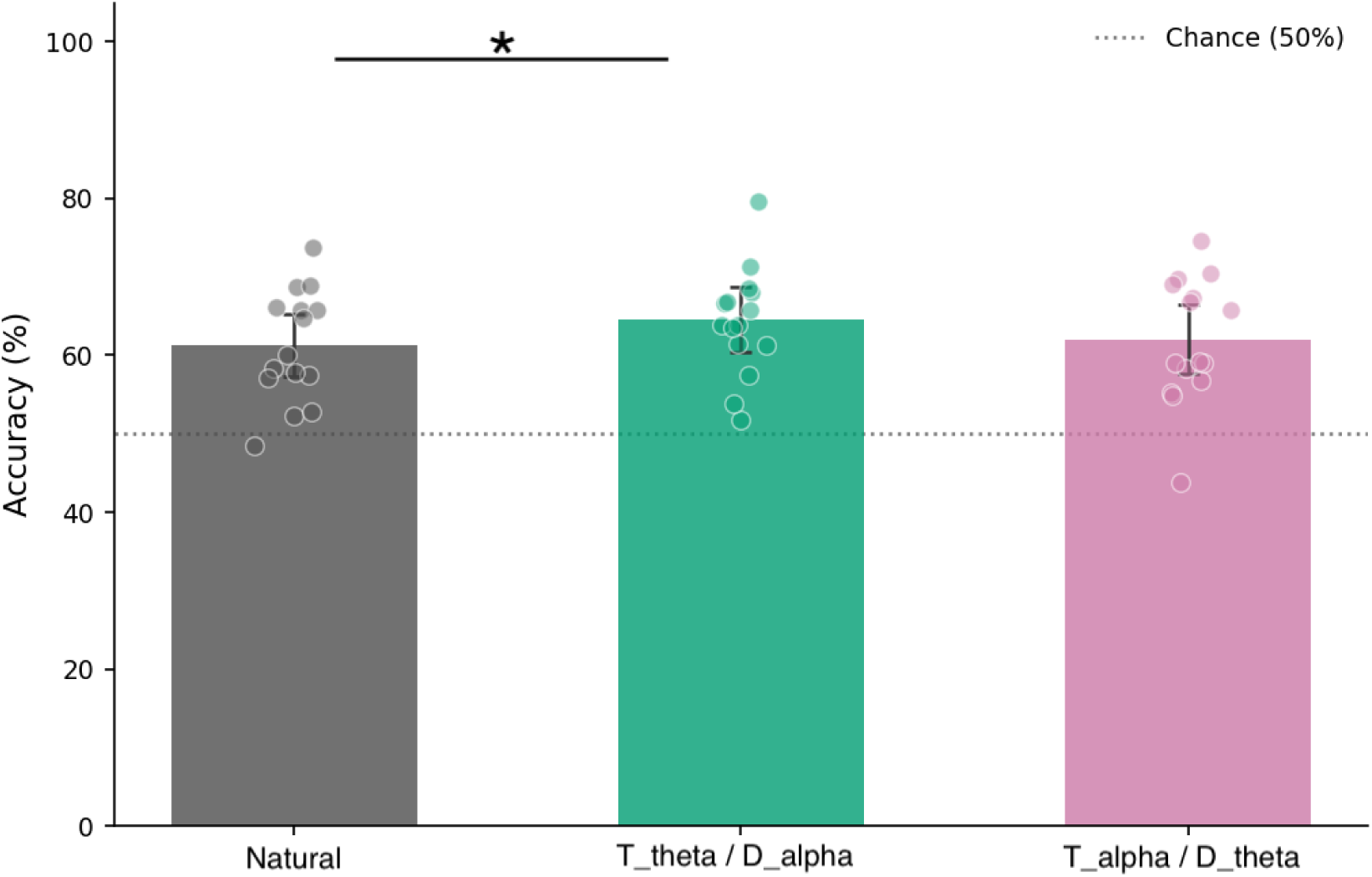
Behavioral performance (accuracy) as a function of block stimulation in the subgroup. Each point represents an individual participant value. Error bars represent 95% confidence intervals. Asterisks indicate significant differences (* *p <* 0.05).

Taken together, these results show that increasing memory load reduced accuracy, whereas informative retro-cues facilitated response speed without producing a significant improvement in accuracy. Rhythmic Gabor flicker did not significantly modulate behavioral performance across the full sample. Exploratory subgroup analyses nevertheless indicated that, among participants who showed no retro-cue benefit in accuracy in the Natural condition at Load 4, accuracy was higher in the *T_theta_/D_alpha_* condition than in the Natural condition, suggesting that this stimulation configuration may be particularly beneficial when baseline prioritization is limited.

### 3.3 Discussion

Experiment 2 tested whether the optimized Gabor flicker stimulation identified in Experiment 1 could be extended to individualized theta- and alpha-frequency stimulation during working-memory retention.

Behaviorally, increasing memory load reduced accuracy, while informative retro-cues produced faster responses than neutral cues. However, retro-cues did not significantly improve accuracy, and rhythmic stimulation did not reliably affect performance across the full sample. One possible contributor to these null group-level effects is the substantial inter-individual variability in cue-based prioritization. In Riddle et al. (2020), participants were behaviorally screened before the stimulation sessions, and those who did not demonstrate a retro-cue benefit were excluded; memory load was also adjusted when necessary to identify a condition in which this benefit was expressed. In contrast, the present study included participants irrespective of their baseline retro-cue benefit and used the same memory loads across individuals. The resulting heterogeneity in the behavioral expression of cue-based prioritization may therefore have reduced sensitivity to stimulation-related changes at the group level and motivated our exploratory examination of baseline-dependent effects.

At the electrophysiological level, however, rhythmic stimulation produced clear frequency-specific responses. Both SNR and ITPC were enhanced in the flicker conditions relative to the Natural condition, demonstrating reliable neural synchronization to individualized theta- and alpha-frequency stimulation during memory retention (Norcia et al., 2015; Vialatte et al., 2010). These findings extend previous work on frequency-specific modulation of working-memory processes by showing that individualized visual stimulation can induce sustained theta- and alpha-frequency responses during an ongoing cognitive task (Riddle et al., 2020).

The contrast between these pronounced electrophysiological effects and the absence of a corresponding improvement in behavior suggests that eliciting frequency-specific neural synchronization is not, by itself, sufficient to enhance working-memory performance. Similar dissociations between neural responses to rhythmic stimulation and behavioral outcomes have been reported previously (De Graaf & Duecker, 2022; Szaszkó et al., 2024). The behavioral consequences of stimulation may therefore depend on additional factors, including task demands and the participant’s baseline cognitive state.

In line with this possibility, the exploratory subgroup analysis suggested that stimulation effects may depend on baseline cue-related performance. Among participants who showed little or no retro-cue benefit in the Natural condition, accuracy was higher when theta-frequency stimulation was applied to cued representations and alpha-frequency stimulation to uncued representations than in the Natural condition. This pattern is compatible with previous reports of greater stimulation-related benefits in individuals with lower baseline working-memory performance or capacity (Pileckyte & Soto-Faraco, 2024; Souza & Oberauer, 2016). Notably, these participants represent a subgroup that would not necessarily have met the behavioral screening criterion used by Riddle et al. (2020), raising the possibility that stimulation responsiveness may differ according to the initial efficiency of cue-based prioritization. Because this analysis was post hoc and based on a reduced sample, however, it should be regarded as hypothesis-generating rather than confirmatory.

Overall, Experiment 2 demonstrates that individualized textured flicker can reliably engage theta- and alpha-frequency neural activity during working-memory retention, while its behavioral consequences appear considerably more variable. These findings emphasize the distinction between neural responsiveness to rhythmic stimulation and measurable cognitive improvement, and suggest that future studies may benefit from prospectively considering baseline cognitive performance, cue responsiveness, and individually calibrated task demands when evaluating behavioral effects of neuromodulation.

## 4 General Discussion

The present study developed and evaluated an individualized rhythmic visual stimulation designed to balance reliable neural entrainment with improved tolerability. Using a stepwise approach, Experiment 1 aimed to identify the stimulation parameters that provided the best balance between neural entrainment and tolerability by manipulating frequency, intensity, and stimulus structure during passive viewing. Experiment 2 then applied the optimized Gabor flicker stimulation during a retro-cue working memory task to assess whether individualized alpha and theta frequency stimulation could modulate behavioral performance and neural activity.

Experiment 1 showed that entrainment depended strongly on the alignment between the stimulation frequency and each participant’s individual alpha frequency (IAF). SSVEP signal quality was highest at IAF and declined when stimulation was shifted 2 Hz below or above this frequency. This pattern is consistent with neural resonance accounts, according to which rhythmic stimulation is most effective when the external driving frequency matches intrinsic oscillatory dynamics (Gulbinaite et al., 2017; Keitel et al., 2019; Notbohm & Herrmann, 2016). Stimulus format also shaped the balance between neural efficacy and tolerability. Although salient flicker can produce strong entrainment, it may also be uncomfortable or intrusive, limiting its use in sustained cognitive tasks (Fisher et al., 2005). In the present study, Plain and Gabor flickers produced comparable overall SNR, whereas Gabor flickers were rated as substantially more comfortable and less mentally fatiguing. Together, these findings identify Gabor stimulation at IAF as the most favorable compromise between entrainment strength and visual comfort.

Experiment 2 extended the frequency-specific neuromodulation framework introduced by Riddle et al. (2020), while replacing transcranial magnetic stimulation with a less invasive, more accessible, and better-tolerated Gabor flicker stimulation optimized in Experiment 1. Beyond the local frequency tagged response, rhythmic visual stimulation may influence cognition by interacting with endogenous oscillations and task-relevant networks. Frequency matching can facilitate phase alignment with intrinsic rhythms (Notbohm & Herrmann, 2016), consistent with the enhanced response at IAF observed in Experiment 1, while rhythmic visual stimulation can modulate activity and connectivity beyond occipital sensory areas (Albouy et al., 2022). In working memory, frontoparietal theta dynamics have been implicated in the prioritization of relevant representations, whereas posterior alpha activity has been associated with the suppression of irrelevant information (Jensen & Mazaheri, 2010; Riddle et al., 2020). Accordingly, theta frequency stimulation of the cued hemifield combined with alpha frequency stimulation of the uncued hemifield could improve working-memory performance by jointly enhancing the prioritization of relevant representations and the suppression of irrelevant ones.

Although individualized theta and alpha frequency stimulation enhanced SNR and ITPC relative to the non-rhythmic Natural condition, reflecting robust frequency specific neural responses and greater phase consistency across trials, these effects did not translate into improved behavioral performance at the group level. These findings highlight a distinction between eliciting reliable frequency specific neural responses and producing measurable cognitive benefits. Similar dissociation between neural entrainment and behavioral outcomes have been reported across cognitive domains (De Graaf & Duecker, 2022; Szaszkó et al., 2024), and theta or alpha frequency visual stimulation has not consistently improved recognition or working memory performance (Wälti et al., 2020; Wimber et al., 2012).

However, the exploratory effect observed in participants with limited baseline retro-cue benefit suggests that individualized rhythmic stimulation may be more effective when baseline cue-based prioritization is limited, leaving greater scope for behavioral improvement. This interpretation is consistent with evidence that the behavioral effects of sensory stimulation can depend on baseline working memory capacity (Pileckyte & Soto-Faraco, 2024; Souza & Oberauer, 2016). Given the exploratory nature and reduced sample size of this analysis, however, this interpretation remains preliminary.

A further limitation concerns the estimation of individualized frequencies from resting state EEG. Although this approach is widely used, resting state peaks may not fully correspond to the oscillatory dynamics expressed during working memory performance, as the amplitude and temporal organization of theta and alpha band activity vary with memory load and task demands (Haegens et al., 2014). Therefore, stimulation frequencies derived from task related oscillatory activity may provide a more precise target for cognitive neuromodulation than resting state estimates alone. In addition, the same memory loads were used for all participants despite substantial differences in baseline performance and the overall difficulty of the task. Future studies should therefore compare resting state and task derived stimulation frequencies and calibrate memory load to each participant’s performance. Such individualization could provide a more comparable level of cognitive demand across participants and help determine whether stimulation effects depend on baseline working memory capacity (Pileckyte & Soto-Faraco, 2024).

Overall, this work provides a stepwise evaluation of individualized Gabor flicker as a tool for visual neuro-modulation. Experiment 1 identified stimulation at the individual alpha frequency as the most favorable balance between SSVEP signal quality and subjective tolerability, while Experiment 2 showed that the optimized stimulation reliably elicited theta and alpha frequency neural responses during working memory retention. However, these electrophysiological effects were not accompanied by consistent behavioral improvements in the full sample, indicating that robust neural entrainment does not necessarily translate into enhanced cognitive performance. Together, these findings support individualized Gabor flicker as a promising and well tolerated approach for probing oscillatory mechanisms during cognition. Future studies should refine stimulation parameters using task-related neural activity, calibrate the memory load to individual performance, and prospectively examine whether baseline working memory capacity predicts behavioral responsiveness.

## Data and Code Availability

The dataset is available for non-commercial scientific research, in accordance with participant consent, ethical approval, and GDPR regulations : https://doi.org/10.57745/UKH1CA. The analysis code is publicly available at the following link: https://doi.org/10.5281/zenodo.23041388.

## Author Contributions

Clément Blanc: Writing – original draft, Methodology, Investigation, Formal analysis, Conceptualization. Frédéric Dehais: Writing review and editing, Validation, Supervision, Project administration, Funding acquisition, Conceptualization. Sébastien Scannella: Writing review and editing, Validation, Supervision, Project administration, Methodology, Conceptualization.

## Declaration of the use of AI

During the preparation of this manuscript, the authors used ChatGPT (OpenAI) to assist with English-language editing and text reformulation. All AI-assisted content was carefully reviewed and revised by the authors.

