## Supplementary data for "Individualized Gabor Flicker for Well-Tolerated Neural Entrainment During Working Memory"

Supplementary Materials

**Supplementary Figure S1.** Visual comfort across flicker conditions.

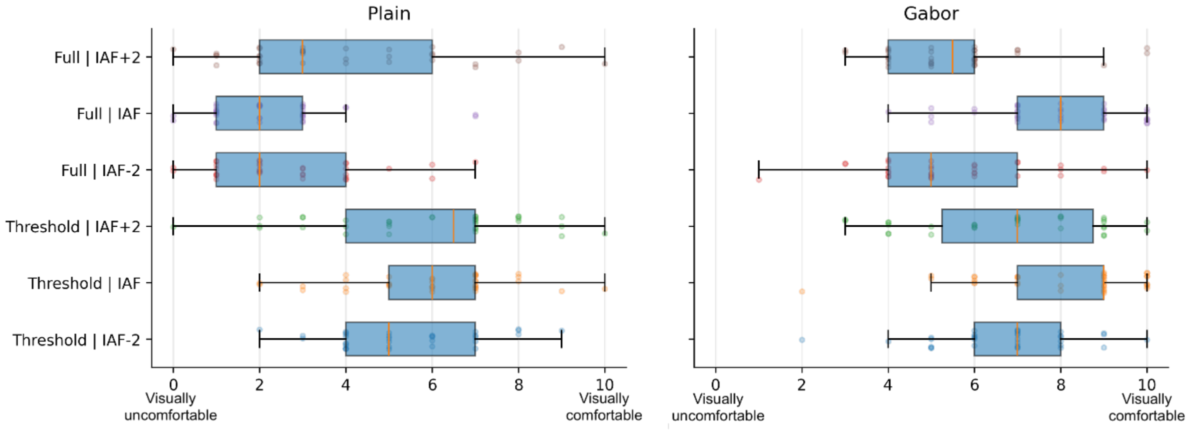


Distribution of subjective visual comfort ratings as a function of flicker intensity (Threshold vs. Full) and stimulation frequency (IAF - 2 Hz, IAF, IAF + 2 Hz), shown separately for Plain and Gabor flicker. Higher values indicate better visual comfort.

**Supplementary Figure S2.** Mental fatigue across flicker conditions.

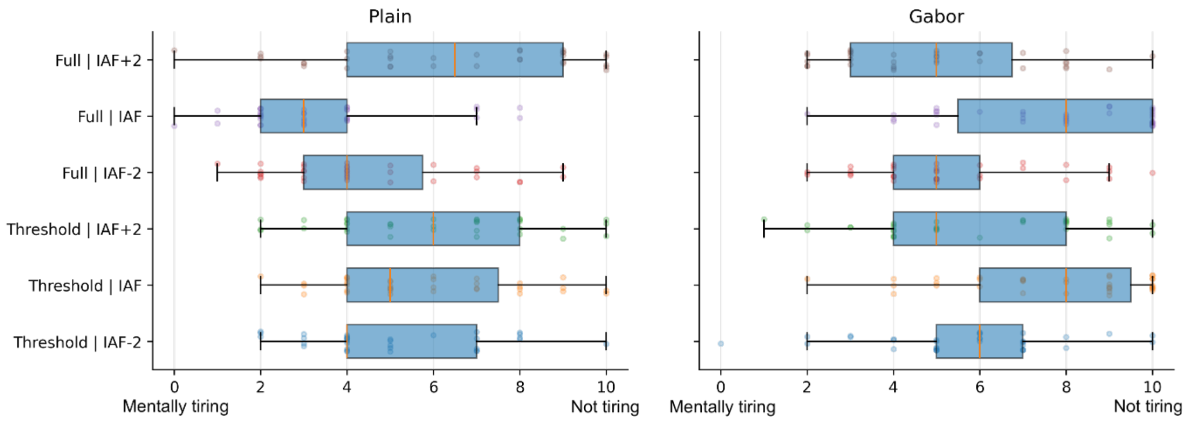


Distribution of subjective mental fatigue ratings as a function of flicker intensity (Threshold vs. Full) and stimulation frequency (IAF - 2 Hz, IAF, IAF + 2 Hz), shown separately for Plain and Gabor flicker. Higher values indicate lower mental fatigue.

**Supplementary Figure S3**. Behavioral performance as a function of cue type.

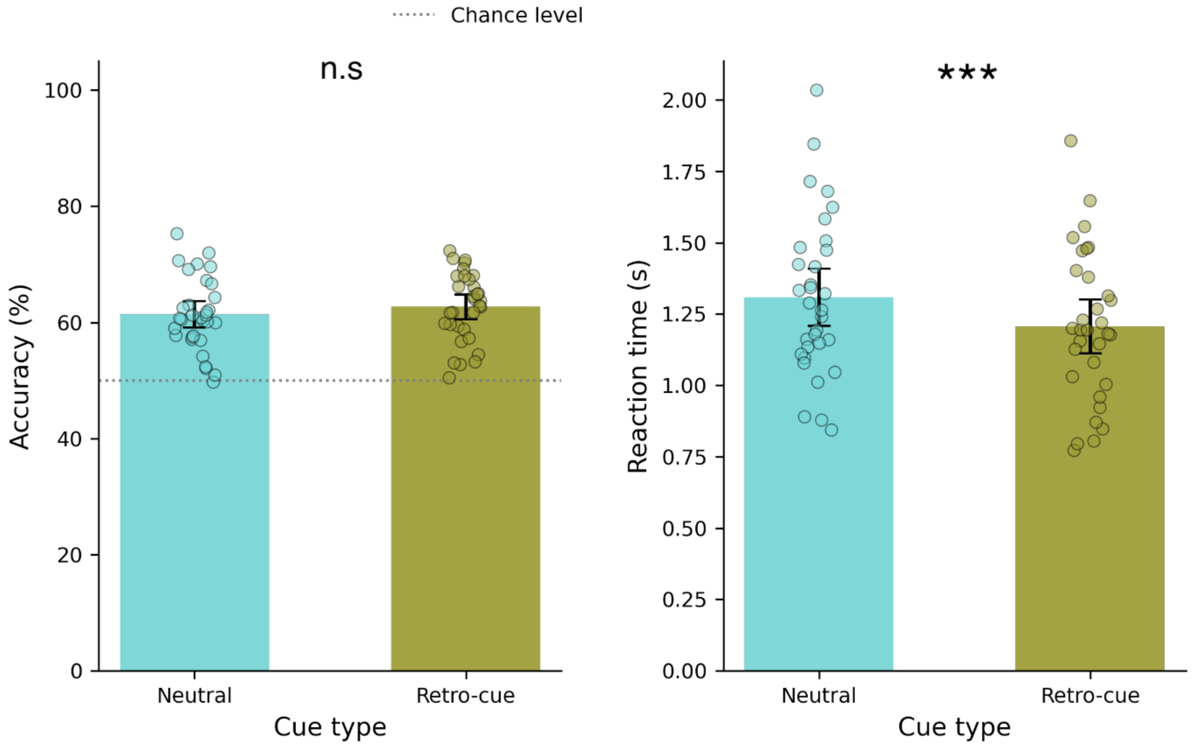


Behavioral performance as a function of cue type (neutral *vs.* retro-cue). The left panel shows accuracy, and the right panel shows reaction times. Each point represents an individual participant value. Error bars represent 95% confidence intervals. Asterisks indicate significant differences (n.s : non significant, *** $p < 0.001$)

**Supplementary Figure S4.** Accuracy across stimulation conditions and memory loads.

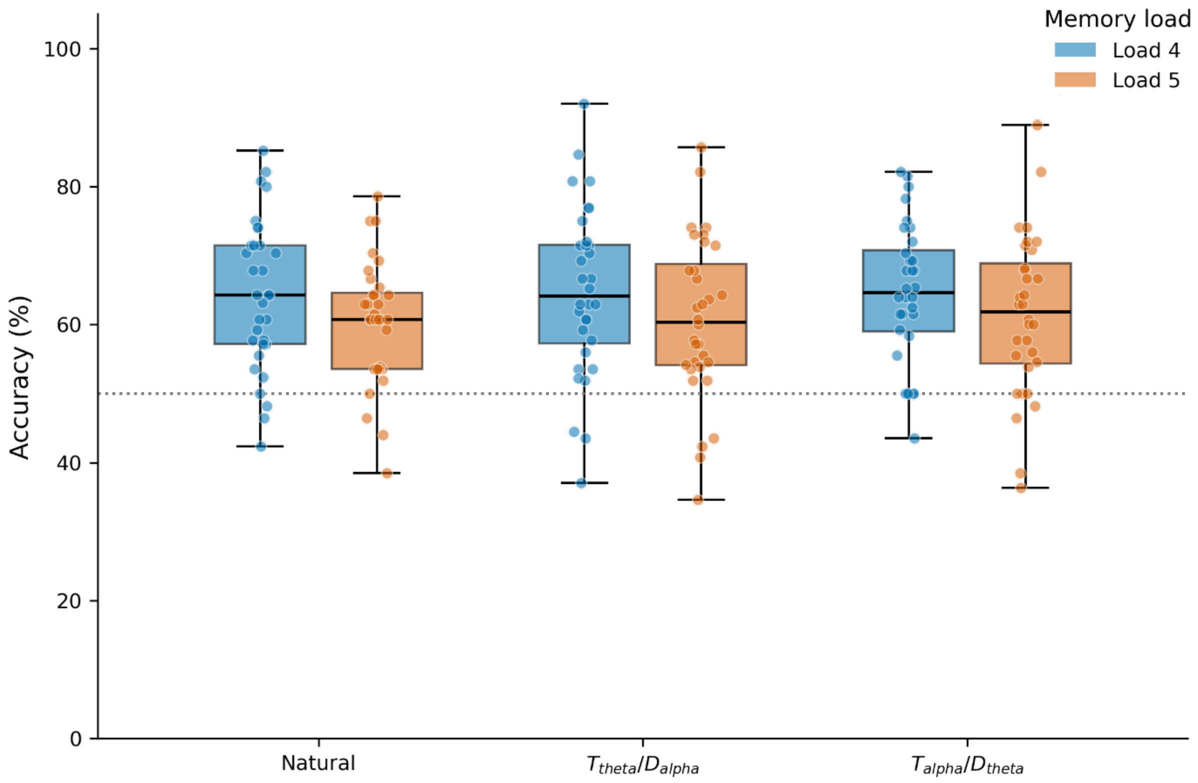


Behavioral performance (accuracy) as a function of stimulation condition and memory load in all participants. Boxplots represent the distribution of individual participant accuracy scores. Accuracy was higher at Load 4 than at Load 5, whereas no main effect of stimulation condition or interaction was observed.

**Supplementary Figure S5.** Reaction time across stimulation conditions and memory loads.

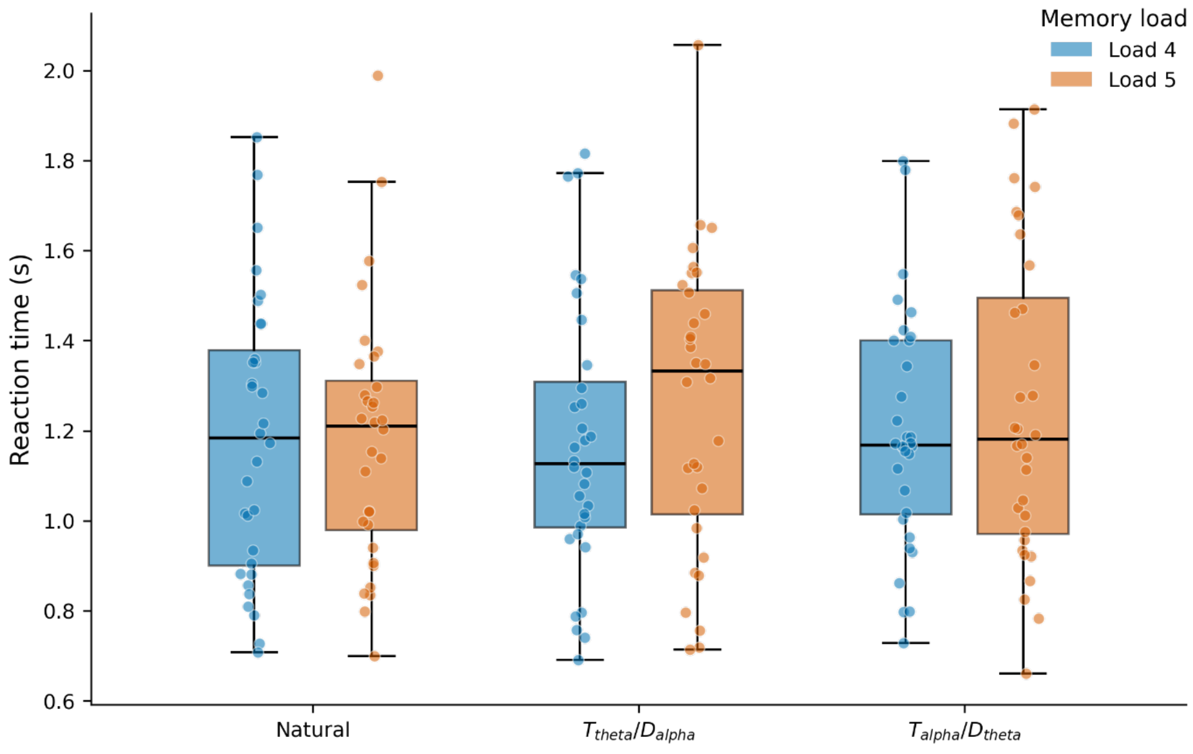


Behavioral performance (reaction time) as a function of stimulation condition and memory load in all participants. Boxplots represent the distribution of individual participant reaction times. Neither stimulation condition nor memory load significantly affected reaction time, and their interaction was not significant.

**Supplementary Video S1.** Example of the experimental paradigm. The video illustrates two example retro-cue trials from Experiment 2, one with memory load 4 and one with memory load 5, in the Target-theta / Distractor-alpha stimulation condition. In both trials, individualized theta frequency flicker was applied to the cued target hemifield during the second retention interval, while individualized alpha frequency flicker was applied to the uncued distractor hemifield.
