## Supplementary figures and images for "Individualized Gabor Flicker for Well-Tolerated Neural Entrainment During Working Memory"

### supplementary_S1_visual_comfort_boxplot.png

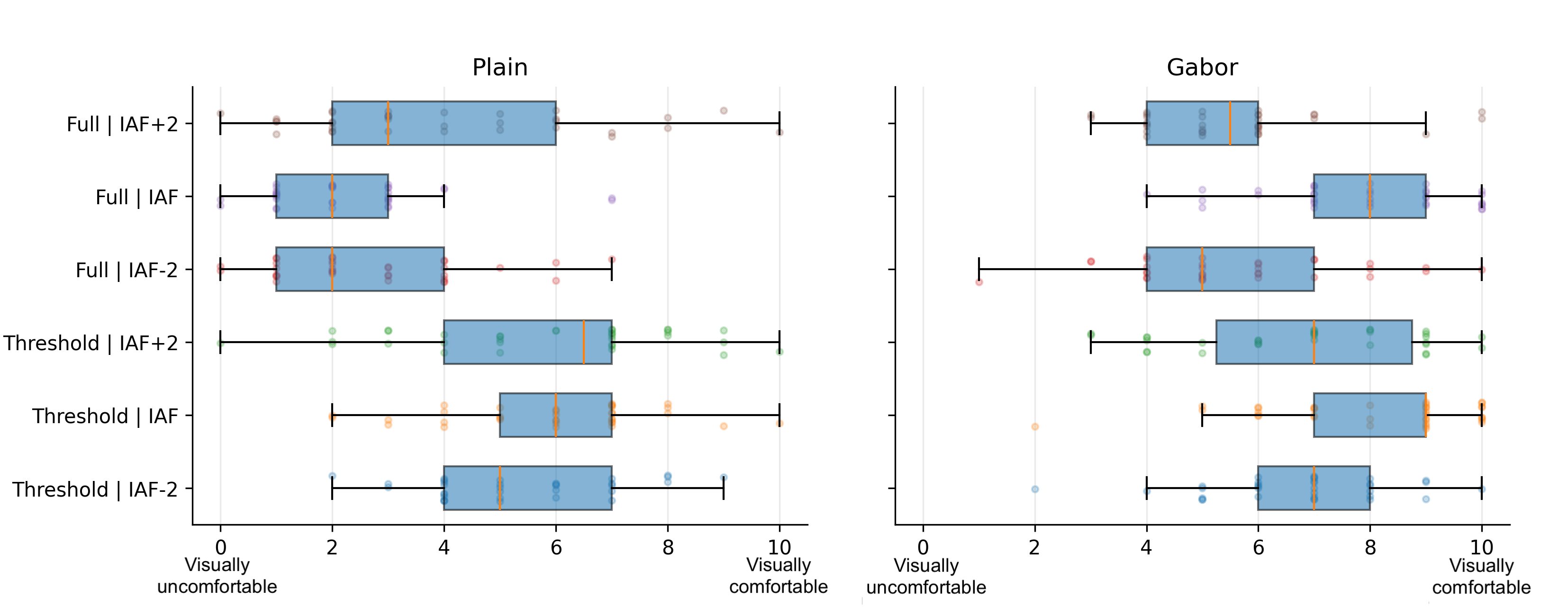

### supplementary_S2_mental_fatigue_boxplot.png

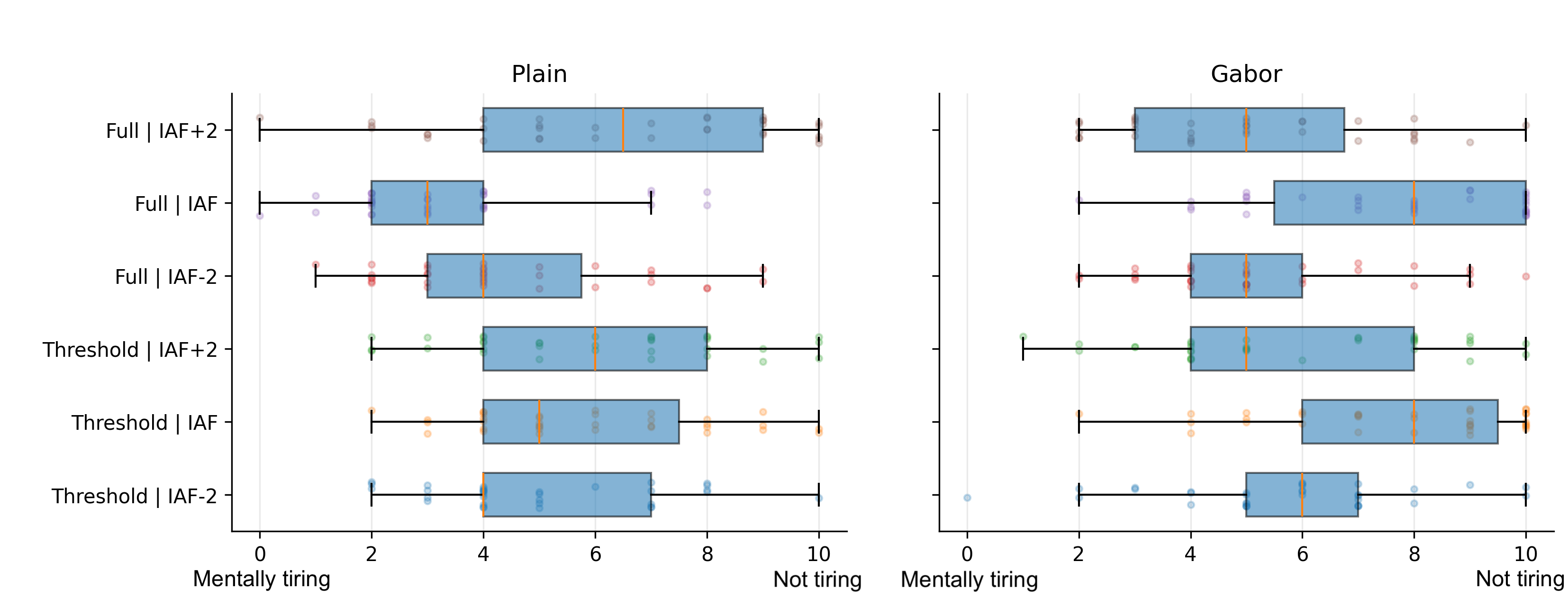

### supplementary_S3_accuracy_rt_by_cue_type.png

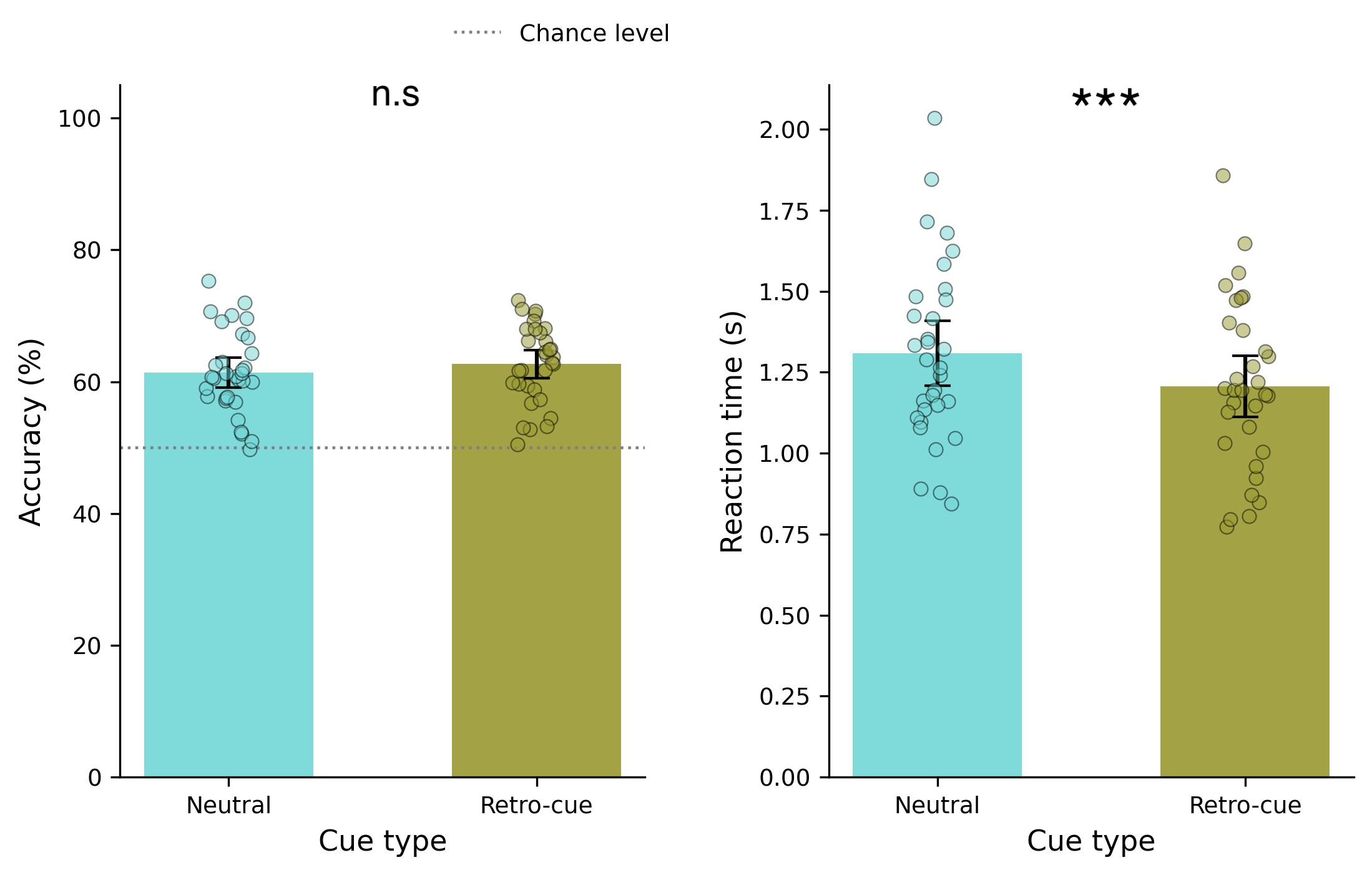

### supplementary_S4_accuracy.png

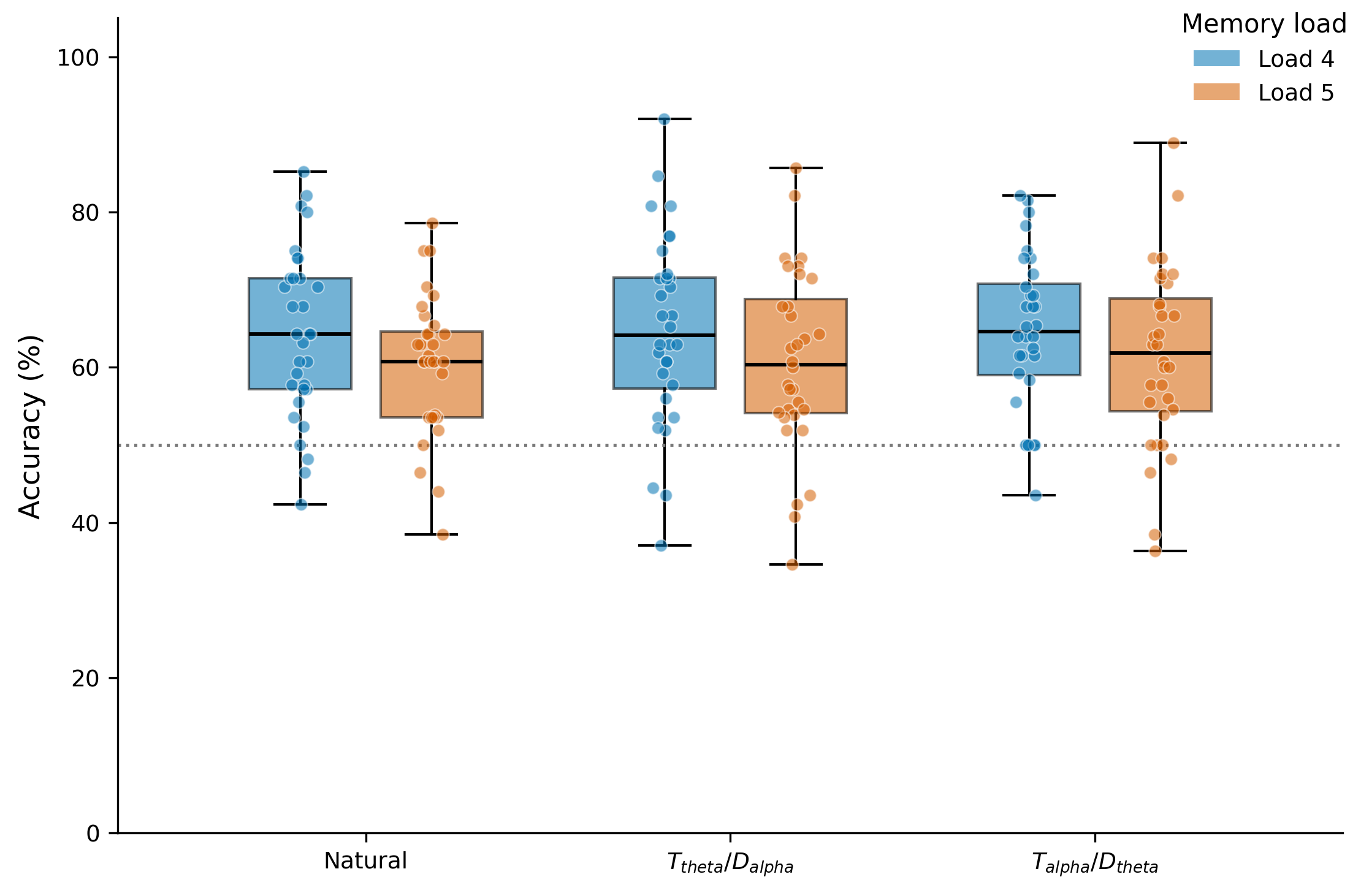

### supplementary_S5_rt.png

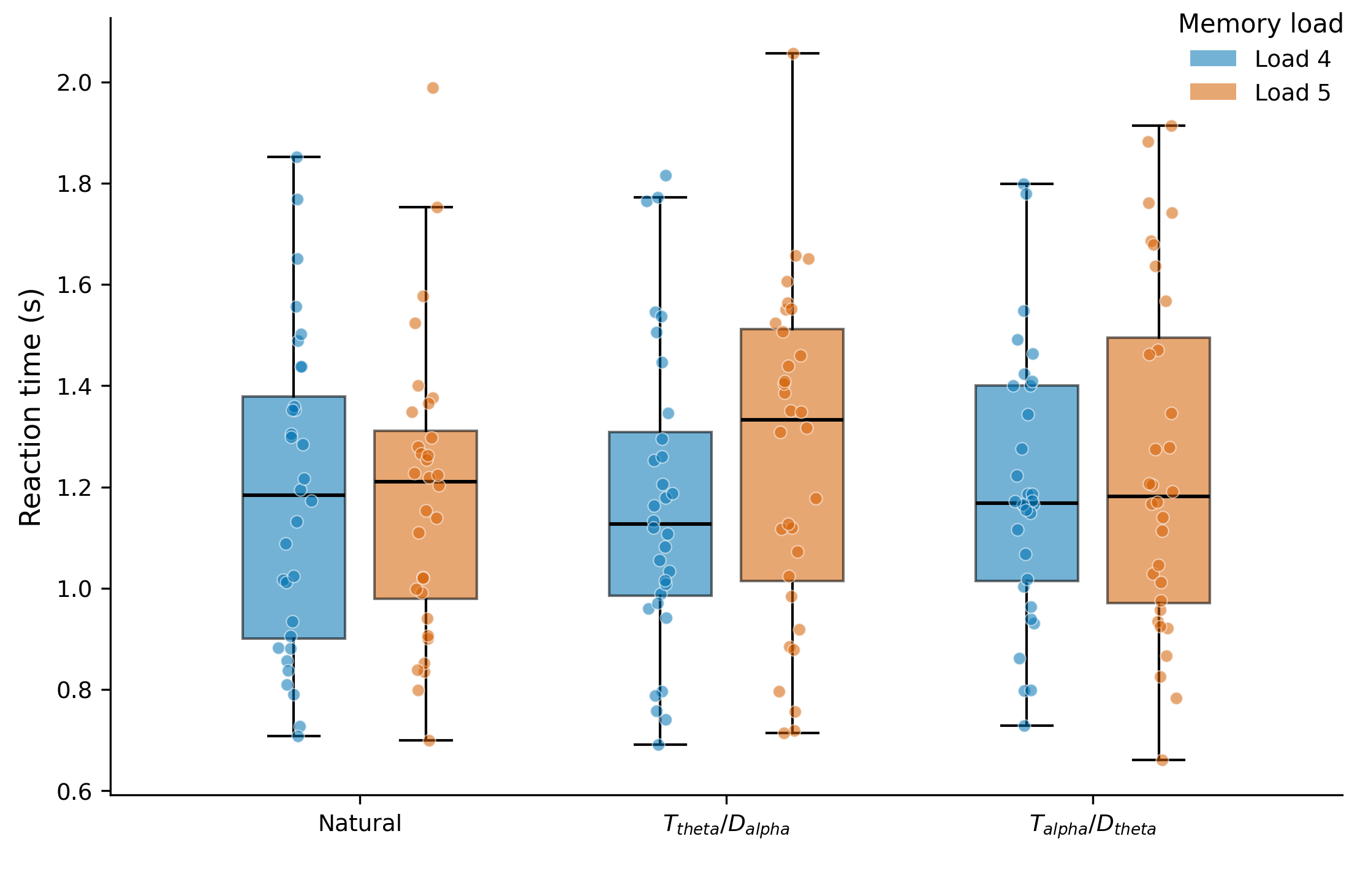
